# Integrated number sense tutoring remediates aberrant neural representations in children with mathematical disabilities

**DOI:** 10.1101/2024.04.09.587577

**Authors:** Yunji Park, Yuan Zhang, Flora Schwartz, Teresa Iuculano, Hyesang Chang, Vinod Menon

**Affiliations:** Department of Psychiatry & Behavioral Sciences, Stanford University, Stanford, CA, 94305; Department of Neurology and Neurological Sciences, Stanford University, Stanford, CA, 94305; Stanford Neuroscience Institute, Stanford University, Stanford, California, CA, 94305; Symbolic Systems Program, Stanford University, Stanford, California, CA, 94305; Centre National de la Recherche Scientifique & Université Paris Sorbonne, Paris 75016, France

**Author notes:** **Address for Correspondence:** Yunji Park, Ph.D. and Vinod Menon, Ph.D. 401 Quarry Rd Stanford University School of Medicine Stanford, CA 94305 *Email:. **Author Contributions:** Y.P., H.C., and V.M. designed research; F.S. and T.I. performed research; Y.P. and Y.Z. analyzed data; Y.P., Y.Z., F.S., H.C., and V.M. wrote the paper. These authors contributed equally to this work (Yunji Park, Yuan Zhang). **Competing Interest Statement:** The authors declare no competing financial interests.

**Keywords:** Neural plasticity, intervention, mathematical disabilities, number sense, multivariate neural pattern analysis

## Abstract

Number sense is essential for early mathematical development but it is compromised in children with mathematical disabilities (MD). Here we investigate the impact of a personalized 4-week Integrated Number Sense (INS) tutoring program aimed at improving the connection between nonsymbolic (sets of objects) and symbolic (Arabic numerals) representations in children with MD. Utilizing neural pattern analysis, we found that INS tutoring not only improved cross-format mapping but also significantly boosted arithmetic fluency in children with MD. Critically, the tutoring normalized previously low levels of cross-format neural representations in these children to pre-tutoring levels observed in typically developing, especially in key brain regions associated with numerical cognition. Moreover, we identified distinct, ‘inverted U-shaped’ neurodevelopmental changes in the MD group, suggesting unique neural plasticity during mathematical skill development. Our findings highlight the effectiveness of targeted INS tutoring for remediating numerical deficits in MD, and offer a foundation for developing evidence-based educational interventions.

**Significance Statement:** Focusing on neural mechanisms, our study advances understanding of how numerical problem-solving can be enhanced in children with mathematical disabilities (MD). We evaluated an integrated number sense tutoring program designed to enhance connections between concrete (e.g. 2 dots) and symbolic (e.g. “2”) numerical representations. Remarkably, the tutoring program not only improved these children’s ability to process numbers similarly across formats but also enhanced their arithmetic skills, indicating transfer of learning to related domains. Importantly, tutoring normalized brain processing patterns in children with MD to resemble those of typically developing peers. These insights highlight the neural bases of successful interventions for MD, offering a foundation for developing targeted educational strategies that could markedly improve learning outcomes for children facing these challenges.

## Introduction

Number sense, the essential capacity to comprehend and manipulate nonsymbolic and symbolic numerical quantities (Berch, 2005; Dehaene, 2011; Jordan et al., 2022; Lau et al., 2021; Locuniak & Jordan, 2008; Mazzocco et al., 2011; Starr et al., 2013), is fundamental for acquiring mathematical skills in early childhood (Devlin et al., 2022; Geary et al., 2013; Holloway & Ansari, 2009; Jordan et al., 2009; Locuniak & Jordan, 2008). Deficiencies in number sense are linked to mathematical disabilities (MD), which affect up to 14% of children (Barbaresi et al., 2005). These disabilities pose considerable hurdles to educational and developmental progress (Butterworth, 2011; Bynner & Parsons, 2005; Rivera-Batiz, 1992). Despite this, many interventions designed to improve number sense are not specifically tailored to meet the needs of children with MD (Bryant et al., 2021; Dyson et al., 2013; Fuchs et al., 2013; Lunardon et al., 2023; Muñez et al., 2022; Obersteiner et al., 2013; Wilson et al., 2006). Furthermore, their effectiveness in enhancing broader mathematical skills remain inconclusive (Lunardon et al., 2023; Maertens et al., 2016; Sella et al., 2016). One leading theory is that MD stems from difficulties in bridging *nonsymbolic* representations (such as sets of objects) with *symbolic* forms (such as Arabic numerals) (Butterworth, 2011; Butterworth et al., 2011; Fias et al., 2013; Kaufmann et al., 2013; Kucian & von Aster, 2015; Price & Ansari, 2013; Rousselle & Noël, 2007). Therefore, interventions focusing on enhancing this cross-format mapping could potentially improve math abilities in children with MD. Despite its importance, there is a notable absence of research on how specific training affects the neural underpinnings of cross-format numerical integration in children with varying levels of math proficiency (see **Table S1** for a summary of prior studies). This gap highlights a critical need for research that explores how targeted training can strengthen the neural connections between different formats of number representation. Addressing this need could lead to more effective, customized interventions for children with MD, ultimately bridging the gap in our comprehensive understanding of number sense development and its impact on mathematical learning.

Here, for the first time, we elucidate the neurocognitive mechanisms underpinning the efficacy of an integrated number sense (INS) tutoring program specifically designed to improve weak number sense in children with MD. Deficits in number sense – the ability to accurately associate symbolic numbers with their corresponding nonsymbolic quantities (Berch, 2005; Jordan et al., 2022; Piazza, 2010) – are a hallmark of MD (Butterworth, 2011; Butterworth et al., 2011; Fias et al., 2013; Kaufmann et al., 2013; Kucian & von Aster, 2015; Price & Ansari, 2013; Rousselle & Noël, 2007). We focus on how children with MD process and map numerical information across these formats. Theoretical models suggest that in typically developing (TD) children, the early stages of learning to associate concrete objects with abstract numerical symbols aid in symbolic numerical problem-solving (Honore & Noel, 2016; Robinson et al., 2002). Interestingly, as TD children gain proficiency, a reverse trend known as “symbolic estrangement” may occur, characterized by distinct neural representations for different number formats (Bulthé et al., 2018; Lyons et al., 2012; Nakai et al., 2023; Schwartz et al., 2021; Wilkey et al., 2020). However, it remains unclear how this typical progression, from the initial integration to subsequent separation of cross-format numerical representations, is altered in children with MD.

Previous investigations have focused on the approximate number system, an innate cognitive system enabling approximate comparisons of nonsymbolic quantities (Geary & vanMarle, 2018; Shusterman et al., 2016; Sullivan & Barner, 2014), as a potential approach for remediating number sense deficits. Despite its intuitive appeal, the efficacy of interventions targeting the approximate number system has been debated, with recent findings suggesting limited impact on enhancing nonsymbolic quantity discrimination or boosting symbolic math skills (Bugden et al., 2021; Kim et al., 2018; Szkudlarek et al., 2021). Moreover, a comprehensive understanding of number sense—which includes the precise alignment of symbolic numbers with their nonsymbolic representations—is increasingly recognized as critical yet deficient in individuals with MD (Butterworth, 2011; Butterworth et al., 2011; Fias et al., 2013; Kaufmann et al., 2013; Kucian & von Aster, 2015; Price & Ansari, 2013; Rousselle & Noël, 2007). Distinct from previous interventions that centered on nonsymbolic number training, the INS tutoring program aims to improve the integration of symbolic number concepts with their nonsymbolic counterparts, addressing a pivotal aspect of number sense not sufficiently covered in earlier studies of children with MD (e.g., Bugden et al., 2021; Szkudlarek et al., 2021).

Our INS tutoring was designed to improve number sense in children with MD, building on previous behavioral and neuroimaging research (Honore & Noel, 2016; Kucian et al., 2011; Lunardon et al., 2023; Michels et al., 2018; Obersteiner et al., 2013; Wilson et al., 2009). This tutoring program consisted of individualized sessions where children progressively learned to associate numbers within and across nonsymbolic and symbolic formats across 4 weeks (**Fig 1A**) (Chang et al., 2022; Chen et al., 2022) We examined behavioral performance and multivariate patterns of brain activity across nonsymbolic and symbolic number comparison tasks, administered to a well-matched sample of 19 children with MD and 21 TD controls during functional magnetic resonance imaging (fMRI) sessions before and after INS tutoring. Additionally, we assessed the broader impact of INS tutoring on standardized arithmetic fluency test performance in these children. This test aimed to determine if the tutoring program could also enhance broader math problem-solving skills in children with MD, which would indicate transfer of learning to other types of mathematical skills that were not directly trained (**Figs 1A-B**).

**Fig 1.**
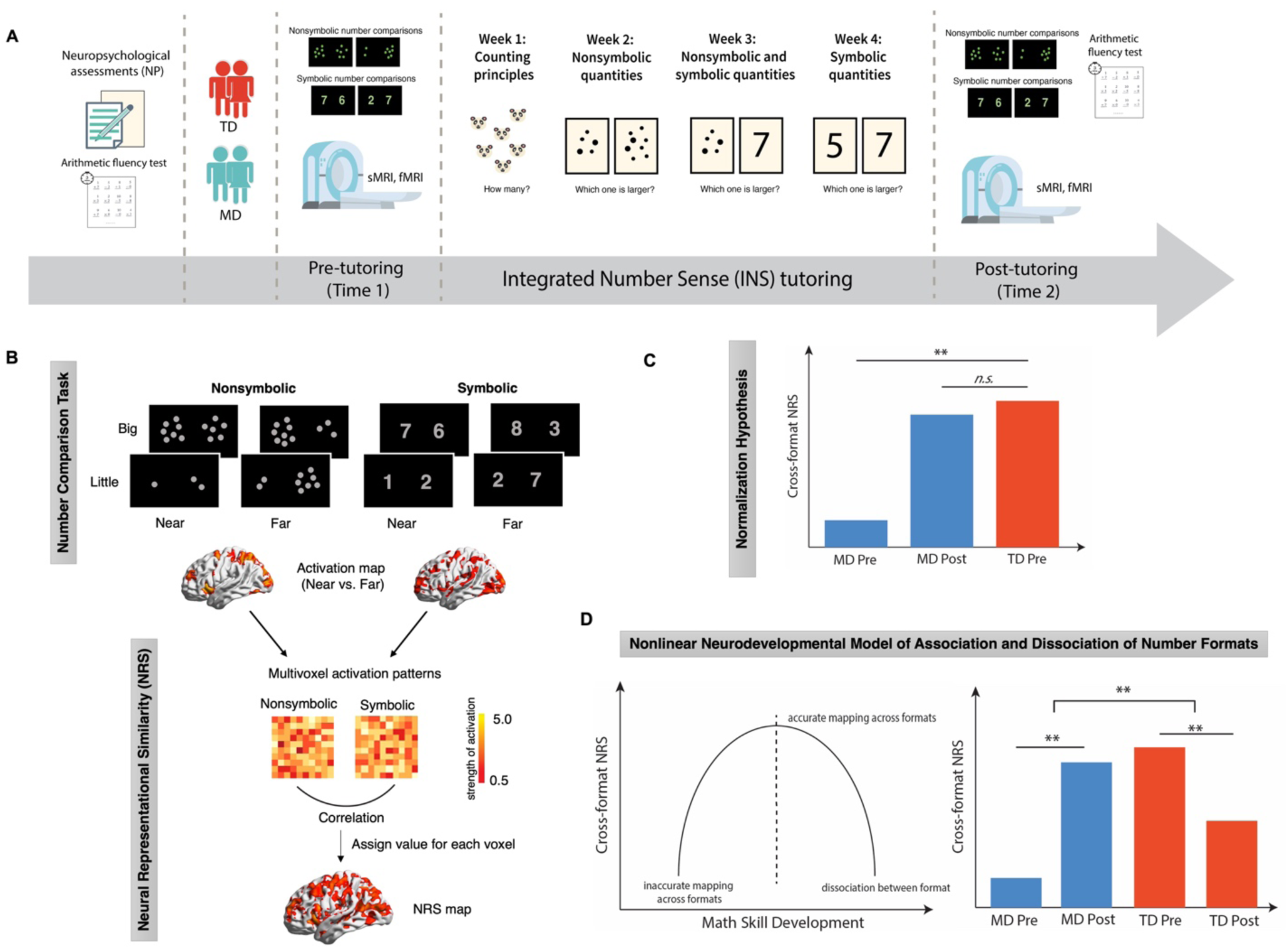
Study design, analysis steps, and key hypotheses. (**A**) ***Study design***. Children with mathematical disabilities (MD) and typically developing (TD) peers underwent pre-tutoring assessments and fMRI scans, followed by a 4-week integrated number sense (INS) tutoring program and post-tutoring assessments and fMRI scans. (**B**) ***fMRI task and analysis***. Participants performed nonsymbolic and symbolic number comparison tasks during fMRI scans. We examined whether INS tutoring induced changes in cross-format neural representational similarity (NRS) between nonsymbolic and symbolic numbers. (**C**) ***Neural normalization hypothesis***. We posited that INS tutoring would enhance cross-format NRS in children with MD at post-tutoring, reaching levels similar to TD peers at pre-tutoring. (**D**) ***Nonlinear neurodevelopmental trajectory of math skill development in MD and TD groups***. Expected outcomes illustrate differential effects of INS tutoring on MD and TD groups, supporting an ‘inverted U-shaped’ nonlinear neurodevelopmental model of cross-format association and dissociation of numbers. **Fig 1B**. was adapted from Schwartz et al. (2021). ** hypothetical significant difference. *n.s.* hypothetical non-significant difference.

Crucially, we employed neural representational similarity (NRS) analysis (Diedrichsen & Kriegeskorte, 2017; Kragel et al., 2018; Poldrack & Farah, 2015; Popal et al., 2019) to probe how training influences the neural mapping between nonsymbolic and symbolic representations of quantity in children with MD. NRS analysis is particularly well-suited for examining how children represent numbers in different formats, a crucial aspect of their mathematical skill development. Cross-format NRS between nonsymbolic and symbolic numbers offers several advantages in this context. First, it allows for precise analysis of the neural mechanisms underlying shared representations across nonsymbolic and symbolic number formats. Understanding such cross-format neural mapping is crucial, as difficulties in integrating the two number formats are often implicated in MD (Butterworth, 2011; Butterworth et al., 2011; Fias et al., 2013; Kaufmann et al., 2013; Kucian & von Aster, 2015; Price & Ansari, 2013; Rousselle & Noël, 2007). Second, cross-format NRS analysis sheds light on brain plasticity in response to intervention designed to integrate the numerical formats. Third, NRS analysis can identify subtle yet significant shifts in neural representations at a fine spatial scale (Haxby et al., 2014; Hebart & Baker, 2018; Kriegeskorte & Kievit, 2013; Kriegeskorte et al., 2008a). This approach has the potential to uncover critical information about associations between multivoxel patterns of brain activity associated with stimuli or tasks beyond the capabilities of single-voxel univariate analyses (Ashkenazi et al., 2012; Chang et al., 2019; Chen et al., 2021; Iuculano et al., 2015; Liu et al., 2023; Sheng et al., 2023). Analysis of cross-sectional data by Schwartz and colleagues has demonstrated developmental variations in neural mapping between the two number formats, showing a clear link in with arithmetic skills (Schwartz et al., 2021). However, no prior study has investigated the impact of training on neural mapping between different number formats in children with MD. To bridge this significant gap, we applied NRS analysis to examine how INS tutoring modifies neural associations between different numerical representations in children with MD, compared to their TD peers.

We had three primary objectives. Our first objective was to evaluate whether the 4-week INS tutoring program could normalize weak levels of similarity in cross-format numerical processing in children with MD closer and bring them to pre-tutoring baseline levels of their TD peers. We hypothesized that INS tutoring would induce a more similar processing across nonsymbolic and symbolic number discrimination in children with MD. Furthermore, we aimed to assess whether the tutoring would also lead to gains in arithmetic fluency. We reasoned that if INS tutoring was effective in facilitating transfer of learning, children with MD would show improvements in broader mathematical problem-solving abilities, reaching the performance of TD children before tutoring.

Our second objective was to determine whether INS tutoring could facilitate *neural normalization* of cross-format similarity in number representations in children with MD. Specifically, we determined whether cross-format NRS between nonsymbolic and symbolic numbers (**Fig 1B**) of children with MD at post-tutoring would align with those of TD children at pre-tutoring. We tested the hypothesis that INS tutoring would normalize cross-format NRS in children with MD, bringing it to the level observed in TD children prior to tutoring (**Fig 1C**).

The third objective of our study was to investigate whether INS tutoring alters cross-format similarity in numerical processing and neural representational patterns in children with MD in a different manner from TD children. Based on the symbolic estrangement account (Bulthé et al., 2018; Nakai et al., 2023; Schwartz et al., 2021), we hypothesized that the patterns of tutoring-induced plasticity in children with MD and TD children would follow a *nonlinear neurodevelopmental model of cross-format association and dissociation of numbers* (**Fig 1D**). Specifically, we predicted that as children with MD improve their ability to map between nonsymbolic and symbolic numbers through INS tutoring, their cross-format similarity in numerical processing and NRS would increase. In contrast, for TD children, who likely achieve accurate cross-format numerical mapping earlier than the MD group, we predicted that the former cohort would exhibit reduced cross-format similarity in numerical processing and NRS following tutoring, which would indicate a shift towards more distinct representations of symbolic numbers. If confirmed, such patterns of findings would delineate distinct, nonlinear trajectories of neurodevelopmental changes in cross-format number representations between children with diverse math abilities.

Our findings advance knowledge of the mechanisms by which INS tutoring facilitates behavioral and neural normalization in children with MD. By testing our hypothesized model, the present study not only highlights the effectiveness of targeted intervention but also provides a window into discovering distinct patterns of learning and neural plasticity across children with diverse cognitive abilities. Importantly, these insights may help guide the development of customized interventions tailored for children with MD.

## Results

### INS tutoring normalizes weak cross-format similarity in numerical processing in children with MD

The first objective of our study was to evaluate the efficacy of a 4-week INS tutoring program in enhancing cross-format similarity in numerical processing in children with MD to the level of their TD peers before tutoring (see **Fig 1A** and **Materials and methods**). As a measure of cross-format similarity in numerical processing across nonsymbolic and symbolic formats, children’s *between-format dissimilarity* was computed as the absolute difference in efficiency scores (accuracy/median RT) between nonsymbolic and symbolic number comparison tasks (|Nonsymbolic – Symbolic|). Lower scores on between-format dissimilarity represented more similar processing of different number formats. At pre-tutoring, the MD group displayed a higher level of between-format dissimilarity – indicative of weak cross-format similarity in numerical processing – compared to the TD group (*t*(41.62) = 2.01, *p* = 0.05, Cohen’s *d* = 0.57) (**Fig 2A**). Similarly, between-format dissimilarity assessed using reaction times was also significantly higher in the MD group compared to the TD group (*p* = 0.020, Cohen’s *d* = −0.70; ***SI Appendix***). However, no significant group difference in between-format dissimilarity was observed when comparing post-tutoring children with MD with pre-tutoring TD children (*ps* > 0.669, Cohen’s *ds <* |0.12|). Additional analyses revealed that these changes in between-format dissimilarity with efficiency scores were largely due to changes in reaction times (***SI Appendix***). Our results demonstrate that INS tutoring effectively reduced discrepancies between nonsymbolic and symbolic number processing in children with MD, with their post-tutoring cross-format similarity in numerical processing aligned with pre-tutoring levels in TD peers.

**Fig 2.**
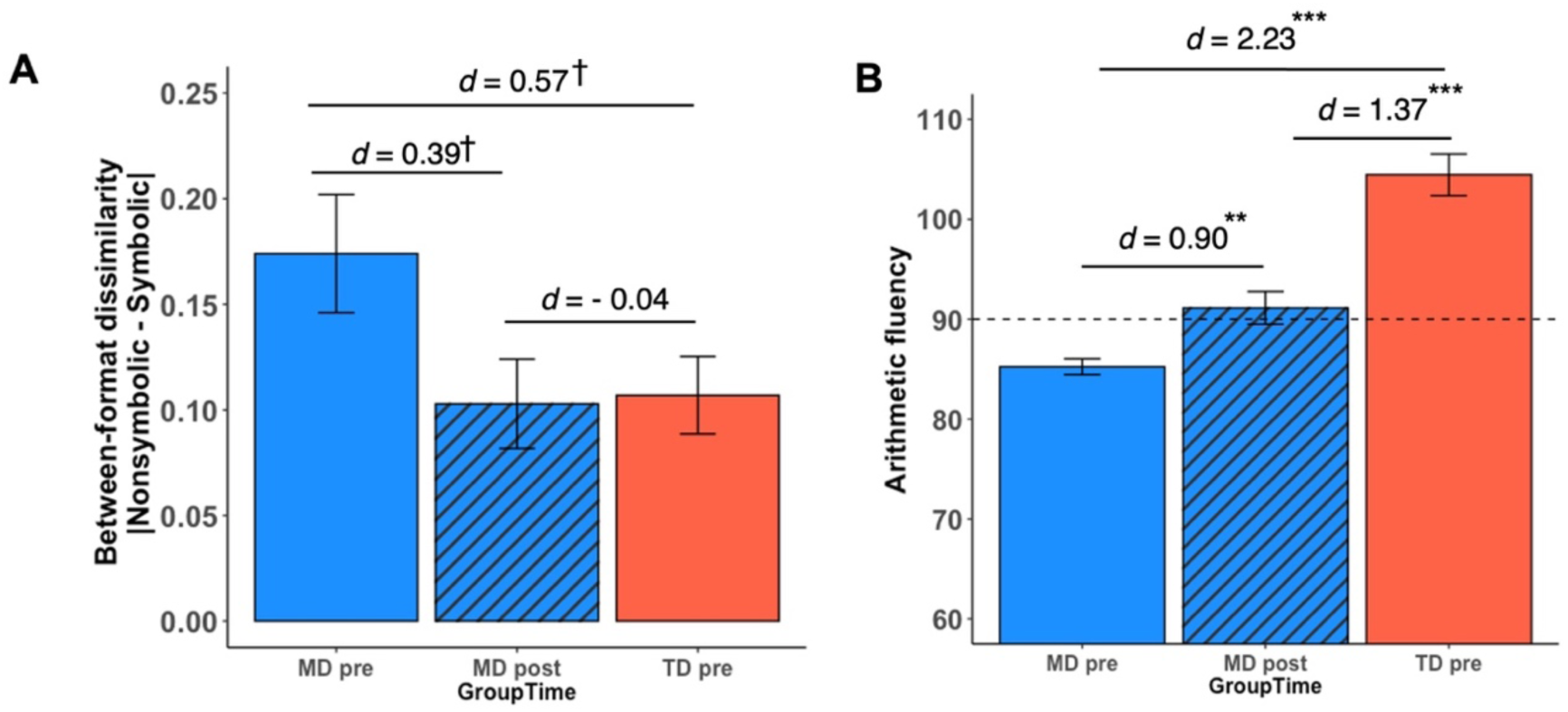
Behavioral normalization of weak cross-format similarity and transfer of learning to arithmetic skills in children with MD following integrated number sense tutoring. (**A**) *Behavioral normalization of weak cross-format similarity between nonsymbolic and symbolic numbers in children with MD*. As a measure of cross-format similarity in numerical processing, a metric of *between-format dissimilarity* was obtained as the absolute difference between nonsymbolic and symbolic number comparison task efficiency. Lower scores on between-format dissimilarity represented higher cross-format similarity in numerical processing. Pre-tutoring, the MD group had higher behavioral between-format dissimilarity (lower cross-format similarity in numerical processing), compared to TD peers. Post-tutoring, this dissimilarity was reduced, aligning with pre-tutoring TD levels. (**B**) *Transfer of learning to arithmetic fluency in children with MD*. Tutoring led to a significant transfer of learning to arithmetic fluency in the MD group, narrowing the pre-existing gap with TD peers. ^†^*p* < 0.10, \*\**p* < 0.01, \*\*\**p* < 0.001, *d* = Cohen’s *d*. Abbreviations: MD, children with mathematical disabilities; TD, typically developing children.

### INS tutoring leads to improvements in arithmetic fluency in children with MD

To further assess the effectiveness of INS tutoring, we examined its impact on arithmetic fluency in children with MD. As the tutoring program specifically focused on enhancing children’s number sense and did not include explicit training of arithmetic skills, children’s gains on arithmetic fluency served as an indicator of transfer of learning to broader math problem solving skills.

At baseline (pre-tutoring), arithmetic fluency in the MD group was significantly lower than that of the TD group (*t*(33.34) = 8.593, *p* < 0.001, Cohen’s *d* = 2.33) (**Fig 2B**). Following the tutoring program, the MD group significantly improved on arithmetic fluency (*t*(25) = 3.41, *p* = 0.002, Cohen’s *d* = 0.90). However, unlike behavioral normalization observed in cross-format similarity in numerical processing, children with MD continued to show lower arithmetic fluency at post-tutoring compared to TD children at pre-tutoring (*t*(48.58) = 5.027, *p* < 0.001, Cohen’s *d* = 1.37) (see **S2 Results** for results from ANOVA). These findings suggest that INS tutoring reduced the gap in arithmetic fluency between the two groups of children, though the performance gap between groups remained following tutoring.

Notably, additional analysis revealed that tutoring-induced gains in arithmetic fluency were significantly related to reduction in between-format dissimilarity in children with MD (*r*(25) = −.404, *p* = 0.045; see details in ***SI Appendix***), which suggests that individual differences in transfer of learning to arithmetic fluency were associated with the degree of changes in cross-format similarity in numerical processing following INS tutoring. These results demonstrate significant improvements in arithmetic fluency in children with MD and suggest that the integration of different number formats may play a role in transfer of learning to arithmetic skills that were not directly targeted by the INS tutoring program.

### Neural normalization in cross-format NRS following INS tutoring in children with MD

Our next objective was to test the neural normalization hypothesis that our INS tutoring leads to normalization of atypical neural representational patterns in the MD group to the level observed in TD children prior to tutoring (**Fig 1C**). To test this hypothesis, we first examined whether cross-format NRS (see **Fig 1B** and **Materials and methods**) are different between children with MLD and TD children before tutoring. We then focused on the identified regions to assess whether post-tutoring cross-format NRS in children with MD became comparable to pre-tutoring cross-format NRS of their TD peers.

At baseline, whole-brain two-sample *t*-test revealed lower cross-format NRS in the MD group compared to the TD group in multiple brain areas, including the hippocampus (Hipp), parahippocampal gyrus (PHG), fusiform gyrus (FG), supramarginal gyrus (SMG), intraparietal sulcus (IPS), and middle frontal gyrus/frontal eye fields (MFG/FEF) (Arsalidou et al., 2018; Sokolowski et al., 2017) (*p* < 0.005, cluster size = 87) (**Fig 3A, Table S3**). Notably, no brain region showed higher cross-format NRS in the MD, compared to the TD, group, which indicates that neural representations of nonsymbolic and symbolic numbers were less similar in the MD group before tutoring.

**Fig 3.**
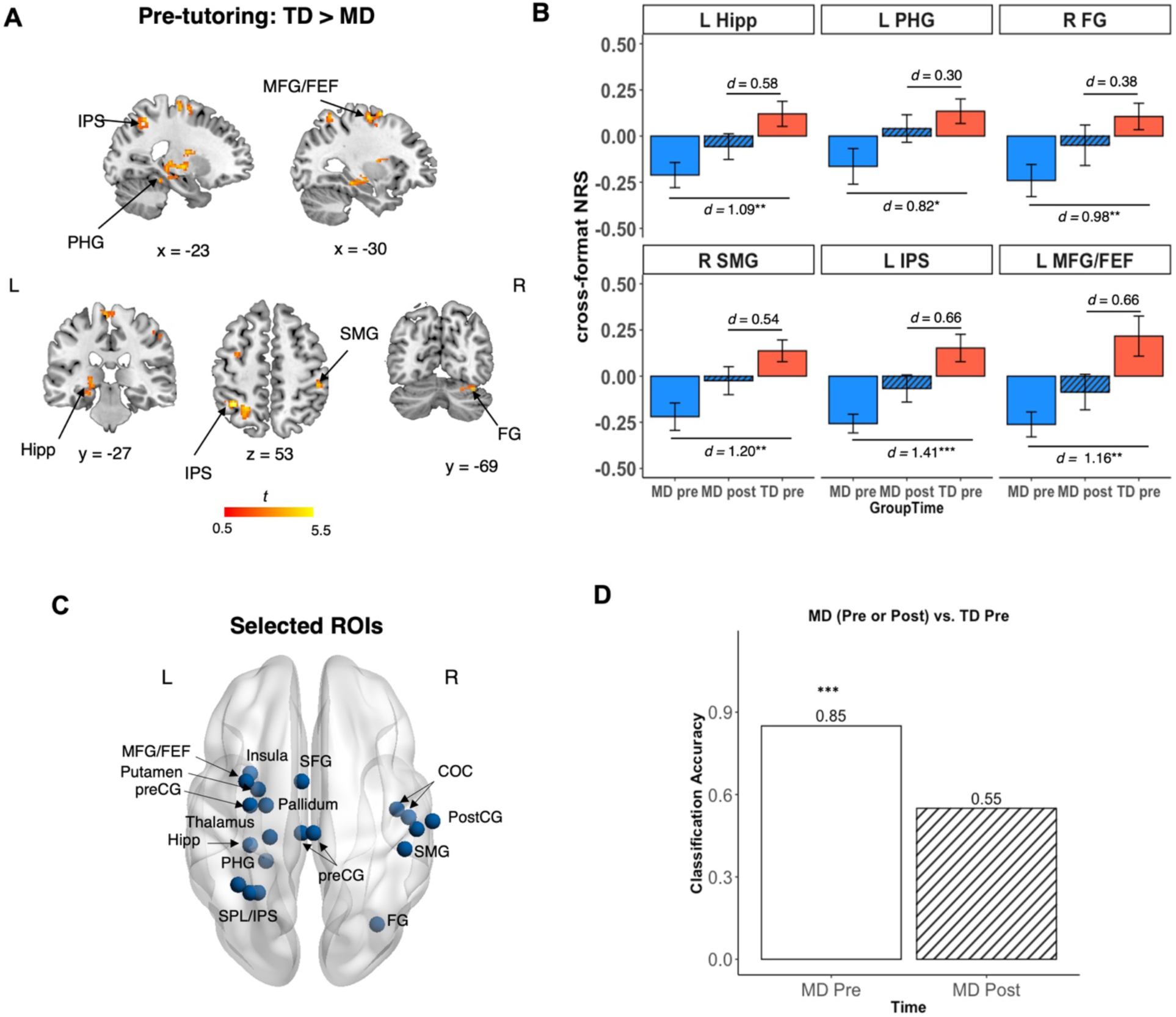
Neural normalization in children with MD following integrated number sense tutoring. (**A**) Whole-brain analysis revealed pre-tutoring differences in cross-format NRS between MD and TD groups in key brain regions implicated in memory and visuospatial processing, including the Hipp, PHG, FG, SMG, IPS, and MFG/FEF. (**B**) Tutoring led to normalization of cross-format NRS in MD at post-tutoring, which was comparable to the pre-tutoring level in TD children (**Table S4**). Effect sizes illustrate a reduction in neural disparity of the MD group following INS tutoring, when compared to the TD group at pre-tutoring, across all brain regions. (**C**) Support-vector machine (SVM)-based classification was performed on 20 brain regions identified from the whole-brain analysis where children with MD exhibited lower pre-tutoring cross-format NRS, compared to their TD peers (**Fig 3A**; see **Tables S3-S4** for details). (**D**) SVM classifier revealed significant difference in cross-format NRS between MD and TD groups at pre-tutoring (*p* < 0.001), but not between MD group at post-tutoring and TD group at pre-tutoring (*p* = 0.36). \**p* < 0.05, \*\**p* < 0.01, \*\*\**p* < 0.001. *d* = Cohen’s *d*. Abbreviations: COC, Central Opercular Cortex; FEF, frontal eye fields; FG, Fusiform Gyrus; Hipp, hippocampus; IPS, intraparietal sulcus; MD, children with mathematical disabilities; MFG, middle frontal gyrus; NRS, neural representational similarity; PHG, parahippocampal gyrus; preCG, precentral gyrus; postCG, Postcentral Gyrus; SFG, superior frontal gyrus; SMG, supramarginal gyrus; SPL, superior parietal lobe; TD, typically developing children. L, Left; R, Right.

Crucially, post-tutoring analyses showed significant increases in cross-format NRS across these regions in children with MD. Among the regions that the MD group had lower cross-format NRS than the TD group before tutoring, we found that INS tutoring led to increases in cross-format NRS in the MD group in all ROIs, reaching levels comparable to the TD group at pre-tutoring (*FDR-corrected p*s > 0.203; **Figure 3B, Table S4**). These results demonstrate that INS tutoring effectively normalized atypical neural patterns in all brain regions observed to be aberrant prior to tutoring in children with MD.

Further validation of the neural normalization hypothesis came from multivariate classification analyses of aberrant NRS in the MD group across distributed brain regions. We used a linear support vector machine (SVM) to conduct a multivariate classification analysis using cross-format NRS from 20 brain regions identified from the whole-brain analysis before tutoring as described above (see **Materials and methods, Table S3, Fig 3C**). SVM significantly differentiated MD and TD groups at pre-tutoring (accuracy = 0.85, *p* = 0.0006; **Fig 4B**) as expected, but failed to differentiate cross-format NRS in the MD group at post-tutoring from that of TD group at pre-tutoring (accuracy = 0.55, *p* = 0.361) (**Fig 3D**). These results underscore that INS tutoring led to network-level neural normalization in the MD group, bringing them on par with TD children at pre-tutoring.

**Fig 4.**
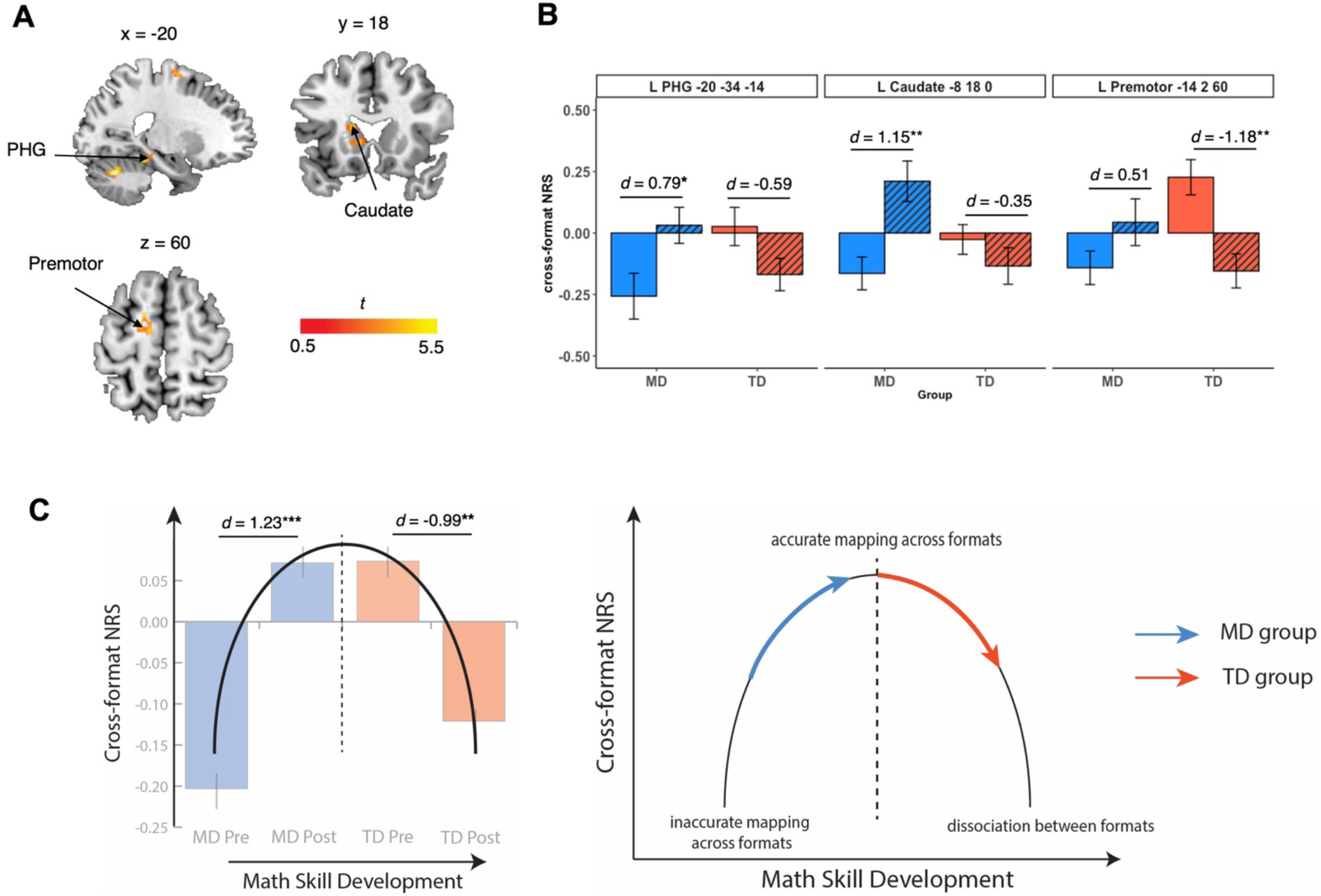
Differential neurodevelopmental trajectories of math skill development in MD and TD groups following integrated number sense tutoring. (**A**) Time (pre-tutoring, post-tutoring) x Group (MD, TD) ANOVA interaction analysis revealed a significant Time x Group interaction on cross-format NRS in 7 brain regions, including the PHG, caudate, and premotor cortex. (**B**) Time x Group interaction effect was characterized by relative increases in cross-format NRS in the MD group and relative decreases in cross-format NRS in the TD group in observed brain regions. (**C**) The observed changes across the two groups aligned with an inverted U-shaped nonlinear neurodevelopmental model of cross-format association and dissociation of numbers. Averaged cross-format NRS across all identified regions from Time x Group interaction effect provided support for distinct learning trajectories between the two groups: children with MD showed enhanced cross-format NRS at post-tutoring, reaching levels similar to pre-tutoring TD levels; TD children shifted towards more specialized, distinguishable neural representations between the two number formats, aligned with their increased proficiency in symbolic numbers. See **Tables S5-6** for details. \**p* < 0.05, \*\**p* < 0.01. *d* = Cohen’s *d*. Abbreviations: MD, children with mathematical disabilities; NRS, neural representational similarity; PHG, parahippocampal gyrus; TD, typically developing children. L, Left; R, Right.

### Nonlinear neurodevelopmental profile of cross-format association and dissociation of numbers

Our final objective was to test a nonlinear neurodevelopmental model of association and dissociation between number formats (**Fig 1D**) by characterizing the patterns of behavioral and neural representational changes in children with MD (i.e., children who are in relatively earlier stages of math skill development) and their TD peers (i.e., children who are in relatively later stages of math skill development) following INS tutoring. We examined whether INS tutoring induced distinct patterns of changes in cross-format similarity in numerical processing (between-format dissimilarity) and NRS in the two groups.

#### Cross-format similarity in numerical processing

In a mixed-design ANOVA with Group (MD, TD) as the between-subject factor and Time (pre-tutoring, post-tutoring) as the within-subject factor, we found a significant interaction between Group and Time on between-format dissimilarity (*F*(1,49) = 10.26 *p* = 0.002, *11*^2^ = 0.08). We found reduced (and increased) between-format dissimilarity in the MD (and TD) group after tutoring, indicative of increased (and decreased) cross-format similarity in numerical processing. No other main effects were significant (*ps* > .840). These findings support our hypothesis that INS tutoring focused on integration of nonsymbolic and symbolic number representations would induce distinct patterns of behavioral changes between children with MD and TD children.

#### Cross-format NRS

To test the neurodevelopmental model of distinct profiles of neural plasticity between children with MD and their TD peers following INS tutoring (**Fig 1D**), we performed a whole-brain ANOVA with Group (MD, TD) and Time (pre-tutoring, post-tutoring) factors on cross-format NRS (see **Materials and methods**, **Fig 4A**, **Table S5**). Here we observed a significant interaction between Group and Time in multiple brain regions, including the PHG, caudate, and premotor area, pointing to differential tutoring-induced neural representational changes in these brain regions between the two groups (see **Fig 4B**, ***SI Appendix*** for details).

Next, we examined the overall profile of neurodevelopmental differences between groups in response to INS tutoring, by averaging cross-format NRS across 7 brain regions that showed a significant Group by Time interaction in the whole-brain ANOVA (**Fig 4C, Table S5**). Here, we found that the significant interaction between Group and Time (*F*(1,76) = 24.95, *p* < 0.001, *11*^2^ = 0.25) was characterized by a significant increase in cross-format NRS in the MD group (*t*(18) = 4.12, *p* < 0.001, Cohen’s *d* = 1.23) and a significant decrease in cross-format NRS in the TD group (*t*(20) = −2.95, *p* = 0.008, Cohen’s *d* = −0.99). These patterns of changes reflect an inverted U-shaped neurodevelopmental model of association and dissociation between two number formats from earlier (MD group) to later (TD group) stages of math skill development (**Fig 4C**).

To further validate these findings at a network level, we conducted a multivariate classification analysis using SVM. We used cross-format NRS data from the 7 ROIs identified as showing significant interaction effects between Group and Time in the whole-brain ANOVA as described above (**Table S5**). SVM accurately differentiated post- vs pre-tutoring differences in cross-format NRS (i.e., neural plasticity patterns) between MD and TD groups (accuracy = 0.80, *p* = 0.001). These results demonstrate distinct patterns of tutoring-induced plasticity in cross-format NRS between the two groups. Together, our findings suggest that INS tutoring induces non-linear patterns of math-ability-dependent changes in neural plasticity between nonsymbolic and symbolic numbers and provide support for the proposed neurodevelopmental model.

## Discussion

We implemented a 4-week, personalized integrated number sense (INS) tutoring program designed to enhance foundational numerical skills in children with mathematical disabilities (MD). Our INS tutoring program targeted the integration of nonsymbolic and symbolic numbers in the earlier stages of tutoring, progressively transitioning to an emphasis on symbolic numerical processing in its later stages. We examined tutoring-induced changes in cross-format similarity in numerical processing and neural representational similarity (NRS) between nonsymbolic and symbolic numbers in children with MD, in comparison to TD controls. A key aspect of our analytic approach was the focus on NRS analysis to characterize how neural connections between nonsymbolic and symbolic number formats evolve with learning. This multivariate neural pattern analysis allowed us to examine if INS tutoring could normalize neural representations of numbers in children with MD, offering a more nuanced understanding than traditional univariate analyses (Chen et al., 2021; Liu et al., 2023; Sheng et al., 2023). By uncovering the underlying neural mechanisms of the remediation of weak number sense in children with MD, our study provides essential insights into effective interventions that address the unique learning needs of these children.

We highlight three key findings, each aligned with specific hypotheses tested. First, INS tutoring remediated weak cross-format similarity between nonsymbolic and symbolic number discrimination in children with MD. Remarkably, INS tutoring also improved arithmetic fluency in children with MD, suggesting that the program could induce transfer of learning to broad math problem-solving skills in these children. Second, multivariate neural pattern analysis revealed normalization of cross-format NRS in children with MD. Post-tutoring, cross-format NRS in children with MD aligned with pre-tutoring level in TD controls, in line with our neural normalization hypothesis. Third, our analysis revealed that INS tutoring induced distinct patterns of change in cross-format similarity in numerical processing and NRS in children with MD compared to their TD peers. Our findings provide evidence for a nonlinear neurodevelopmental model that describes the shift from integration to segregation of nonsymbolic and symbolic number representations across different stages of mathematical skill development. These findings deepen our understanding of learning and neural plasticity in children with MD and have implications for the development of personalized intervention for children with neurodiverse abilities.

### 4 weeks of INS tutoring remediates weak cross-format similarity in numerical processing in children with MD to the level of TD controls

The first objective of our study was to determine the effectiveness of a 4-week INS tutoring in remediating behavioral measure of cross-format numerical mapping in children with MD compared to TD children. We examined whether the two groups of children show different behavioral profiles of similar processing across nonsymbolic and symbolic numbers before tutoring and whether INS tutoring can normalize atypical levels of similar cross-format number processing in children with MD. Prior to tutoring, children with MD showed lower cross-format similarity between nonsymbolic and symbolic number comparison, compared to TD children.

This result aligns with previous research suggesting that children with MD have less precise mapping and different processing ability between symbolic numbers and nonsymbolic quantities (Butterworth, 2011; Butterworth et al., 2011; Fias et al., 2013; Kaufmann et al., 2013; Kucian & von Aster, 2015; Price & Ansari, 2013). Remarkably, following tutoring, children with MD exhibited a level of cross-format similarity in numerical processing that is comparable to their TD peers prior to tutoring. Thus, our results suggest that the 4-week INS tutoring program was effective in remediating low levels of similarity in processing across nonsymbolic and symbolic numbers in children with MD.

We further assessed transfer of learning to broader math problem-solving skills in children with MD by examining the impact of the INS tutoring program on their arithmetic fluency. Notably, children with MD showed significant improvements on arithmetic fluency even though they were specifically trained on basic number sense, such as counting and comparison, and were not instructed on arithmetic principles. Unlike other interventions that incorporate explicit instructions on arithmetic problem solving (Kucian et al., 2011; Michels et al., 2018; Tobia et al., 2021), our program was specifically designed to bolster number sense across different formats. This finding is particularly noteworthy given the scarcity of intervention studies focusing on symbolic and nonsymbolic cross-format numerical mapping without introducing arithmetic in children with MD. Our results suggest that targeted number sense training can contribute to broader gains in arithmetic problem-solving skills for children with MD even in the absence of explicit arithmetic training.

Additionally, our analysis revealed a significant relation between improvements in arithmetic fluency and changes in cross-format similarity in numerical processing in children with MD indicating that better integration between nonsymbolic and symbolic number formats can directly bolster arithmetic skills. This connection emphasizes the critical contribution of number sense to broader mathematical competencies. Together, our findings highlight the effectiveness of cross-format number sense training in not only promoting learning, but also meaningful transfer to arithmetic problem-solving, specifically in children with MD.

### Neural normalization of cross-format number representations in children with MD

The second objective of our study was to test the neural normalization hypothesis by examining whether aberrant levels of cross-format NRS can be remediated by INS tutoring in children with MD. At pre-tutoring baseline, the results revealed lower cross-format NRS in the MD group compared to the TD group in distributed brain areas spanning the medial temporal lobe and ventrotemporal occipital, parietal, and prefrontal cortices. This included the hippocampus, parahippocampal gyrus, fusiform gyrus, intraparietal sulcus, supramarginal gyrus, and middle frontal gyrus/frontal eye field. Crucially, after INS tutoring, levels of cross-format NRS in the MD cohort across all these brain regions became comparable to those of TD children at their baseline, which indicates that tutoring normalized aberrant neural representations in children with MD. Furthermore, multivariate classification analysis revealed a lack of distinction between post-tutoring cross-format NRS in children with MD and pre-tutoring cross-format NRS in TD children. These findings provide converging evidence that the INS tutoring program facilitated neural representational plasticity in children with MD by strengthening neural mapping between nonsymbolic and symbolic numbers to baseline levels observed in their TD counterparts.

Brain regions that displayed atypical cross-format NRS patterns in children with MD before tutoring are known to be critical for numerical cognition (Ansari, 2016; Geary & Moore, 2016; Hyde & Ansari, 2018; Iuculano et al., 2018; Menon & Chang, 2021). The hippocampus and parahippocampal gyrus are crucial for integration of relational memories and the binding of disparate information into cohesive cognitive structures (Eichenbaum, 2004; Giovanello et al., 2004; Olsen et al., 2012; Ranganath, 2010; Staresina & Davachi, 2009), a process likely essential for forming associations between nonsymbolic and symbolic number formats (Menon & Chang, 2021). The fusiform gyrus plays a crucial role in representing complex visual symbols including numbers (Arsalidou & Taylor, 2011; Chen et al., 2021; Hannagan et al., 2015; Iuculano et al., 2018; Skagenholt et al., 2022). The intraparietal sulcus is associated with representation and manipulation of numerical quantity as well as spatial attention (Ansari, 2016; Cantlon et al., 2006; Holloway & Ansari, 2010; Hubbard et al., 2005; Kersey & Cantlon, 2017; Menon & Chang, 2021; Sokolowski et al., 2017). The middle frontal gyrus and supramarginal gyrus are involved in higher-level cognitive functions such as task-switching or working memory (Aron et al., 2004; Barbey et al., 2013; Deschamps et al., 2014; Li et al., 2022; Silk et al., 2010). It is noteworthy that INS tutoring led to normalization of aberrant cross-format NRS in children with MD in all these brain regions, reaching baseline levels in TD children.

Taken together, our findings indicate that the neural remediation of cross-format number representations in children with MD involves a network of distributed brain areas, which aligns with systems neuroscience models of neural systems and pathways that are implicated in impairments underlying MD (Iuculano et al., 2018; Menon & Chang, 2022; Menon et al., 2020). Our findings suggest that INS tutoring had a substantial impact on these interconnected neural systems to remediate MD. This neural normalization in children with MD demonstrates the efficacy of INS tutoring and emphasizes its potential in enhancing the conceptual association between different numerical formats.

### Divergent changes in cross-format numerical processing and neural plasticity in children with MD and TD children following INS tutoring

The third objective of our study aimed to determine whether the INS tutoring program differentially influences behavioral and neural plasticity in children with MD and TD children. We tested a nonlinear neurodevelopmental model of cross-format association and dissociation of numbers across varying stages of mathematical skill development. This model was informed by a theoretical account of ‘symbolic estrangement,’ which posits a developmental trajectory where children gradually dissociate symbolic numbers from their corresponding nonsymbolic quantities as they gain proficiency in numerical skills (Bulthé et al., 2014; Lyons et al., 2012; Schwartz et al., 2021).

Our analysis revealed significant Time x Group interactions for both behavioral and neural measures of cross-format numerical mapping, pointing to diverging trajectories of changes in cross-format numerical processing and neural representational plasticity following tutoring. Behaviorally, children with MD showed a marked increase in cross-format similarity in processing across nonsymbolic and symbolic numbers, while TD children, in contrast, showed a decrease in this cross-format similarity after tutoring. In line with our hypothesis, these contrasting behavioral profiles suggest that INS tutoring exerts distinct, ability-specific effects on cross-format numerical mapping depending on children’s initial math ability.

At the neural level, our NRS analysis revealed a significant Time x Group interaction in multiple brain areas, including the parahippocampal gyrus, caudate, and premotor area. This interaction effect was characterized by an inverted U-shaped curve of cross-format NRS between nonsymbolic and symbolic numbers across math skill development, aligned with our hypothesized nonlinear neurodevelopmental model (**Fig 4**). Specifically, the MD group, who initially exhibited a lower cross-format NRS indicative of suboptimal neural mapping, showed an increase in cross-format NRS post-tutoring, which suggests enhanced neural mapping following tutoring in this cohort. In contrast, TD children, who initially had higher cross-format NRS at baseline, exhibited a decrease in cross-format NRS post-tutoring, indicating a shift towards more efficient symbolic numerical processing in these children.

These divergent patterns of behavioral and neural plasticity between children with MD and TD children corroborate our proposed nonlinear neurodevelopmental model and provide new insights into the varied impact of INS tutoring depending on the stage of math skill development in each group. These findings underscore the importance of accounting for initial math ability when aiming to mitigate learning disparities between children with MD and their TD peers and optimizing learning for all children. Our findings advance our understanding of neurodevelopmental pathways in numerical cognition, offering valuable insights for tailoring effective, personalized educational interventions.

## Conclusion

Our study sheds light on the neural mechanisms through which a short-term integrated number sense tutoring program effectively remediates weak number sense in children with MD. We further discovered that children with MD display distinct tutoring-induced learning and neural plasticity pathways when compared to their typically developing peers. In a significant advance over univariate analyses, our application of multivariate neural pattern analysis yielded deeper insights into how the brain organizes and differentiates numbers in different formats and how tutoring influences neural plasticity across varying mathematical abilities.

Our integrated number sense tutoring program not only improved number sense but also arithmetic problem-solving abilities in children with MD, indicating transfer of learning to broader math problem-solving skills that are critical for positive educational outcomes. Importantly, tutoring normalized aberrant neural representational patterns in children with MD to align with those of typically developing children at baseline. This normalization occurred across distributed brain regions, indicating that tutoring induced widespread functional reorganization in the brains of children with MD. Furthermore, our results highlight nonlinear neurodevelopmental trajectories of neural representational changes that occur across varying levels of mathematical skill development, significantly advancing our understanding of diverse neural pathways across various stages of learning in children.

Our findings are relevant for developing evidence-based pedagogical strategies to close the performance gap between children with MD and their typically developing peers. These insights underscore the importance of designing tailored interventions that address the unique learning needs of children with MD, including the integration of various numerical formats to enhance their learning outcomes. The development of effective interventions should take into consideration of neurodiverse trajectories to further improve educational experiences of children facing learning challenges.

## Methods and Materials

### Participants

We recruited a total of 66 children in the second or third grade of elementary school from multiple school districts. The protocol was approved by the Institutional Review Board and informed written consent was obtained from the child’s legal guardian. The research has been conducted in accordance with institutional ethical guidelines and the Declaration of Helsinki.

For behavioral data analysis, a total of 13 participants were excluded due to unmatched cognitive abilities between TD and MD groups (*N* = 2), low birth weight (*N* = 1), missing data (*N* = 5), task administration error (*N* = 2), below chance performance in both comparison tasks (*N* = 2) and brain abnormality (*N* = 1). A total of 53 children were included in behavioral data analysis (27 female, 25 children with MD, *Meanage* = 8.16, *SDage* = 0.66).

Additional 13 participants were excluded from fMRI data analysis due to excessive head movement (*N* = 9), poor image quality (*N* = 3), and poor co-registration of brain images to a template (*N* = 1). A total of 40 children were included in fMRI data analysis (22 female, 19 children with MD, *Meanage* = 8.18, *SDage* = 0.58).

MD and TD groups were defined using criterion-based cutoff scores from standardized math fluency test. Children with MD scored at or below 90 (i.e., the 25^th^ percentile) (e.g., Fuchs et al., 2004; Wilson & Swanson, 2001) and TD children scored above 90 on Math Fluency subtest of the Woodcock Johnson-III (WJ-III) (Woodcock et al., 2001). The two groups of children were not significantly different on age, gender, and general cognitive abilities (see also ***SI Appendix*** and **Table S2** for details).

### Experimental Procedures

The present study aimed to examine children’s cognitive and brain plasticity in response to INS tutoring. Children underwent cognitive assessments and brain imaging sessions before and after a 4-week INS tutoring. The INS intervention included 1-on-1 tutoring sessions. Detailed task descriptions are below (see also **Fig 1A**).

#### Nonsymbolic and symbolic number comparison tasks

Before and after tutoring, children completed nonsymbolic and symbolic number comparison tasks in the scanner wherein they determined the larger between two nonsymbolic (dot arrays) or symbolic (Arabic numerals) numbers presented on each side of the screen (**Fig 1B**). A total of 64 trials was presented in each run. Numbers between 1 and 9, excluding 5, were presented. We used a 2 × 2 experimental design accounting for both the size of the number pair (little, big) and the distance between the numbers of the pair (near, far), resulting in 16 trials per condition. In half of the trials, the sum of the pair was greater than 10 (“big” numbers), and in the other half, it was less than 10 (“little” numbers). In half of the trials, the distance between the numbers was 1 unit (“near” distance), and in the other half, the distance was 5 units (“far” distance). Additional details on number comparison tasks are described in ***SI Appendix***.

#### Cognitive Assessments

Children completed a battery of cognitive assessments to assess their math abilities as well as IQ, reading abilities, and working memory (see details in ***SI Appendix***). Demographics and scores from cognitive assessments at the time of inclusion are shown in **Table S2**.

#### Arithmetic Fluency

Children’s math ability was assessed from Math Fluency subtest of the WJ-III (Woodcock et al., 2001). This subtest is a timed pencil and paper test that measures individual’s ability to quickly and accurately solve simple addition, subtraction and multiplication problems (**Fig 1A**). Children were given a 3-minute time limit and instructed to solve as many problems as they can. All problems were presented vertically and involved operands from 0-10.

#### INS Tutoring

Across four weeks, children underwent twelve sessions (3 sessions/week) of one-on-one tutoring specifically designed to enhance fundamental understanding of relations between nonsymbolic and symbolic numbers, as well as each number format processing. Quantities ranged from 1 through 9 to facilitate exact processing of numbers. Learning activities for children progressed gradually each week, aiming to build proficiency in exact symbolic number processing: children learned and practiced basic counting principles in *week 1*, processing of nonsymblic quantities in *week 2*, understanding relations between nonsymbolic and symbolic quantities in *week 3*, and lastly, processing of exact symbolic numbers in *week 4*. In each session, a trained tutor used various interactive learning tools including physical manipulatives and computer games. At the end of each session (except for first two sessions), children completed a review worksheet, which included a list of problems based on the week’s focus. Children received stickers upon completion of activities. Additional details on the tutoring protocol are described in **SI.**

### Statistical analysis

### Behavioral analysis

#### Number comparison tasks

For nonsymbolic and symbolic number comparison tasks, trials with response times lower than 150ms were excluded from the analysis. We computed children’s efficiency scores by dividing accuracy by median reaction times (RT) for each task to account for potential speed-accuracy trade-off (Townsend & Ashby, 1978). To aid interpretation of the results, we opted to use efficiency score (Edwards et al., 2015; Hoffman & Schraw, 2009), instead of inverse efficiency score (IES) (MacLeod & Nelson, 1984). Dissimilarity between nonsymbolic and symbolic number processing (between-format dissimilarity) was assessed by calculating absolute difference in efficiency between two comparison tasks. For this analysis, two children’s data were excluded due to poor performance in one (nonsymobolic or symbolic) comparison task.

To test *behavioral normalization hypothesis*, planned two-sample *t*-tests assessed (i) whether children with MD exhibit higher between-format dissimilarity than TD children before tutoring and (ii) whether between-format dissimilarity in children with MD after tutoring reach the level of TD children before tutoring. To further understand whether tutoring induced similar or different patterns of changes in between-format dissimilarity in number comparison ability between children with MD and TD children, we performed a mixed ANOVA with Group (MD, TD) as the between-subject factor and Time (pre-, post-tutoring) as the within-subject factor on between-format dissimilarity. Follow up two-sample and paired *t*-tests clarified significant effects from ANOVA.

#### Arithmetic Fluency

As part of behavioral normalization hypothesis, planned two-sample *t*-tests assessed (i) group difference in arithmetic fluency (measured by WJ-III Math Fluency) before tutoring and (ii) difference in arithmetic fluency between MD group after tutoring and TD group before tutoring. To further understand tutoring-induced changes in arithmetic fluency in the two groups of children, we performed a mixed ANOVA with Group (MD, TD) as the between-subject factor and Time (pre-, post-tutoring) as the within-subject factor on arithmetic fluency. Follow up two-sample and paired t-tests clarified significant effects from ANOVA.

To report effect size for all behavioral analyses, we used a generalized eta squared (*h*^2^) for ANOVA (Bakeman, 2005) and Cohen’s *d* for *t*-tests (Cohen, 1992). Absolute Cohen’s d values of 0.2, 0.5, and 0.8 indicate small, medium, and large effect sizes, respectively.

### fMRI data analyses

#### fMRI data acquisition and preprocessing

Functional images were acquired on a 3T GE scanner (General Electric, Milwaukee, WI) using an 8-channel GE head coil. A T1-weighted, 132 slice high-resolution structural image was acquired at both pre- and post-tutoring scan sessions to facilitate registering each participant’s data to standard space. Head movement was minimized using additional pads and pillows around children’s head. A T2*-sensitive gradient echo spiral in-out pulse sequence (Glover & Lai, 1998) was acquired with the following parameters: repetition time (TR) = 2000ms, echo time (TE) = 30ms, flip angle = 80°, field of view (FOV) = 220 mm, matrix size = 64 × 64, resolution = 3.44 × 3.44 × 4.5 mm^3^, interleaved. A total of 31 axial slices were acquired, 4 mm in thickness and 0.5mm in spacing, covering the whole brain.

Images were preprocessed and analyzed using SPM12 (https://www.fil.ion.ucl.ac.uk/spm/). The first five volumes of each time-series were discarded to allow for signal equilibration. The preprocessing pipeline included realignment, slice-timing correction, coregistration to subjects’ T1 and normalization to a 2mm adult Montreal Neurological Institute (MNI) template, similar to a standard practice in neuroimaging research with children (Cantlon et al., 2006; Kersey & Cantlon, 2017; Nakai et al., 2023; Perrachione et al., 2016), and smoothing using a 6mm full-width half-maximum Gaussian kernel to decrease spatial noise. Transitional (*x*,*y*,*z*) and rotational (pitch, roll, yaw) movement parameters were generated from the realignment procedure. Subjects included in the fMRI analysis completed both pre- and post-tutoring scan sessions with following criteria: 1) translational and rotational movement in any direction was less than 12mm and 2) mean scan-to-scan displacement did not exceed 0.7 mm.

#### First-level statistical analysis

Task-related brain activation was assessed using the general linear model (GLM) implemented in SPM12. At the individual subject level, brain responses representing correct trials for each condition (i.e., little near, little far, big near, big far) were modeled using boxcar functions of 2500ms corresponding to the length of a trial convolved with a canonical hemodynamic response function and a temporal derivative to account for voxel-wise latency differences in hemodynamic response. An error regressor was also included in the model to account for incorrect trials. Additionally, both transitional and rotational head movement parameters were included as regressors of no interest. Serial correlations were accounted for by modeling the fMRI time series as a first-degree autoregressive process. The GLM was applied to nonsymbolic and symbolic number comparison tasks separately. Voxel-wise contrast maps were generated for each participant for each task. The contrast of interest was the Near vs Far, which corresponds to the neural distance effect, to assess neural representations of quantity while controlling for low-level stimulus features and response demands.

#### Neural representational similarity (NRS) analysis

To assess similarity in neural representation of quantity between formats, spatial correlation of multivariate patterns of brain activity between nonsymbolic and symbolic number comparison tasks was computed across the whole brain for each individual and each session. In contrast to measuring brain activation levels, NRS analysis provides a way to assess whether cognitive processes share similar neural features and to determine which brain areas are most sensitive to overlapping neural representation across nonsymbolic and symbolic number comparisons (Kriegeskorte et al., 2006b; Kriegeskorte et al., 2008b). The NRS approach is based on well-grounded theories and findings on population coding and distributed representations (Kragel et al., 2018) and is ideal for examining underlying representations of mental states or cognitive functions.

Using a searchlight mapping method (Kriegeskorte et al., 2006a), we obtained cross-format NRS of Near vs Far contrast across nonsymbolic and symbolic formats in the neighborhood surrounding each voxel of each individual’s brain. Specifically, a 6-mm spherical region centered on each voxel was selected, and cross-format similarity was computed within the sphere using the spatial correlation of voxel-wise brain activation (beta-weights). Searchlight maps were then created for every individual by going through every voxel across the whole brain. These searchlight maps were subsequently used for second-level analyses.

A few sets of second-level analyses were then conducted. First, to test the *neural normalization hypothesis*, we performed a whole-brain two-sample *t*-test contrasting MD and TD groups at pre-tutoring to identify the regions where the MD group shows atypical cross-format NRS levels compared to those of TD peers. Cross-format NRS of identified regions was compared between the MD group at post-tutoring and the TD group at pre-tutoring. Next, to test the nonlinear *neurodevelopmental model of cross-format association and dissociation of numbers* and identify significant interaction between groups, we performed a whole-brain mixed ANOVA with Group (MD, TD) as a between-subject factor and Time (Pre, Post) as a within-subject factor.

Subsequent whole brain *t*-test contrasting pre-tutoring vs. post-tutoring was performed to confirm the findings of ANOVA (see ***SI Appendix***). All statistical maps were masked with a grey matter mask, and significant clusters were identified using a height threshold of *p* < 0.005, similar to current practices in the fields of developmental cognitive neuroscience (Kersey et al., 2019; Krinzinger et al., 2011; Matejko & Ansari, 2019; Schwartz et al., 2021), with whole-brain family-wise error rate correction at *p* < 0.01 (spatial extent of 87 voxels) based on Monte Carlo simulations. Follow-up regional-level analysis was based on estimated cross-format NRS in the regions identified from whole-brain analysis. False discovery rate (FDR) correction was used to correct for multiple comparisons in regional-level analysis.

#### Multivariate classification analysis

To further confirm our results, we employed multivariate classification analysis using cross-format NRS values. Our classification analysis allowed us to test the neural normalization hypothesis and nonlinear neurodevelopmental model of cross-format association and dissociation of numbers at a larger scale across multiple brain regions.

First, to test neural normalization hypothesis, we used 20 regions of interest (ROIs) identified from the whole-brain two-sample *t*-test contrasting MD and TD groups at pre-tutoring, which included parietal and frontal regions, including the intraparietal sulcus (IPS), superior parietal lobule (SPL), middle frontal gyrus/frontal eye fields (MFG/FEF), superior frontal gyrus (SFG), supramarginal gyrus (SMG), precentral (preCG) and postcentral gyrus (postCG), the hippocampus (Hipp) and parahippocampal gyrus (PHG) in the medial temporal lobe (MTL), the caudate, putamen and pallidum in the basal ganglia, the insula, and fusiform gyrus (FG) (see details in **Table S2**, **Fig 3C**). We investigated whether cross-format NRS in the brain regions showing deficits in the MD group compared to their TD peers at pre-tutoring could be normalized post-tutoring, such that cross-format NRS in the MD group at post-tutoring is non-distinguishable from that of TD group at pre-tutoring. Thus, we performed a classification analysis to distinguish MD group at pre- or post-tutoring from TD group at pre-tutoring.

Second, to test the nonlinear neurodevelopmental model of association of number formats, we used 7 ROIs identified from the whole-brain ANOVA interaction between Time (pre, post) and Group (MD, TD) analysis (see details in **Tables S5-6, Fig 4A**). We investigated whether MD and TD groups show distinct patterns of changes in response to tutoring by performing a classification analysis to examine whether the MD group was distinguishable from the TD group based on differences in cross-format NRS between pre- and post-tutoring (Post – Pre). Thus, we performed a classification analysis to confirm whether cross-format NRS showed the patterns of neural changes in response to training were significantly different between MD and TD groups.

For all classification analyses, a linear support vector machine (SVM) classification algorithm with 10-fold cross-validation, was used to assess the discriminability of cross-format NRS across each ROI set between groups. The python scikit-learn package (https://scikit-learn.org/) was used to perform this analysis. Permutation test (*n* = 5000) was used to assess the significance of classification accuracy.

## Acknowledgements

This research was supported by grants from the National Institutes of Health (HD094623, HD059205, MH084164) and National Science Foundation (DRL-2024856) to V.M. and Stanford Maternal & Child Health Research Institute Postdoctoral Support Award to H.C. We thank participating families and Miriam Rosenberg-Lee, Shelby Karraker, Sarit Ashkenazi, Lang Chen, Kaustubh Supekar, Dietsje Jolles, Dawlat El-Said, Emma Adair, Samantha Mitsven, Laxman Dhulipala, and Sangeetha Santhanam for assistance with the study. We also thank Kristen Pilner Blair for assistance with the development of the Restaurant Game as part of number sense training program. Generative AI software was not used during writing process of this manuscript.

