## Supplementary material for "Integrated number sense tutoring remediates aberrant neural representations in children with mathematical disabilities": SI Appendix

### Methods and Materials

#### Definition for MD and TD groups

MD and TD groups were defined using criterion-based cutoff scores from math tests similar to previous studies investigating children with MD (1-6). Children with MD scored at or below 90 (i.e., the 25th percentile) and TD children scored above 90 on Math Fluency subtest of the Woodcock Johnson-III (WJ-III) (7). We acknowledge that we do not make a distinction between children with persistent developmental dyscalculia and those with milder forms of math disabilities as we did not test if these children had math difficulties that persisted for at least 6 months. Importantly, we identified children with MD and TD children based on specific performance differences in math scores and comparable scores on other domain-general cognitive measures (see S2 Table).

#### Nonsymbolic and Symbolic Number Comparison fMRI tasks

On each trial, a fixation appeared for 500ms followed by a pair of quantities which remained visible for 1000ms and a blank screen for 1500ms to fill up the response phase. Stimuli were presented for short duration to avoid counting of dots in the nonsymbolic condition. For the nonsymbolic number comparison task, (i) the total area covered by each array of dots and (ii) the average size of dots in each array were controlled to account for potential confounds with number of items. Each pair of quantities was presented 4 times, twice with the larger number on the left, and twice with the larger number on the right.

Using a button box, children indicated which quantity was larger by pressing the left button if the larger number was on the left side or the right button if the larger number was on the right side. Stimuli were presented using E-Prime and displayed using an LCD projector and a back-projection screen in the scanner suite. The inter-trial interval between trials was jittered randomly between 1.7 and 3.8 seconds. Total run duration was ~6 min.

#### Cognitive Assessment

*IQ.* The Wechsler Abbreviated Scale of Intelligence™ (WASI; Wechsler, 1999), was administered to measure Verbal (VIQ), Performance (PIQ), and Full-Scale (FSIQ) IQ.

*Reading Abilities.* Reading abilities were assessed using Letter Word Identification and Word Attack subtests of the WJ-III (Woodcock et al., 2001). In the Letter Word Identification subtest, children were asked to fluently read letters and words of increasing difficulty. In the Word Attack subtest, children were asked to read pseudo words of increasing complexity.

*Working Memory.* Eight subtests of Automated Working Memory Assessments (AWMA) were used to assess working memory ability. Digit Recall and Word Recall subtests assessed verbal short-term memory. Backward Digit Recall subtest assessed verbal working memory. Block Recall subtest assessed visuo-spatial short-term memory. Spatial Recall subtest assessed visuo-spatial working memory.

#### INS Tutoring Protocol

*Week 1.* In Week 1, children were introduced to the counting principle through lessons and a video clip demonstrating accurate or inaccurate counting of sock puppets. Children also played Restaurant Game (8, 9), in which they counted the number of dishes to cook for animals presented in the screen. At the end of Week 1, children completed a review worksheet, in which they counted the number of animals on the worksheet. After completion of Week 1 sessions, children were asked to verbally count from 1 to 9 before beginning sessions in Weeks 2-4.

*Week 2.* In Week 2, children were introduced to comparison of nonsymbolic numbers using sets of erasers. Children played Math Circles wherein they determined a Math Circle with more erasers between two circles presented on the table. This lesson from week 2 was designed to increase children's familiarity with number comparison. Starting from week 2, two interactive games with a tutor were introduced to children: 1) Math War (10) in which children compared which of two quantities is larger and 2) Comparing speed in which children determined the quantity of one value above or below the given quantity. In the Math War, the child and the tutor had a deck of card with nonsymbolic numbers for each, and both flipped the card one at a time. The child wrote down the number on his/her own card and the one on the tutor's card, and then determined the card with larger number. In the Comparing speed, the tutor laid four cards with nonsymbolic quantities from 4 to 7 on the table and then the child and tutor each took five cards from their own deck of cards. Next, they placed the card from their own deck on the top of the card on the table if the quantity on the card from their deck was below or above the quantity of the card on the table. When the child placed all the cards from their own deck, the game was completed. Lastly, children completed a review worksheet for Week 2, in which they determined which of two nonsymbolic numbers was larger.

*Week 3.* In Week 3, children were introduced to integration of nonsymbolic and symbolic representations of numbers through similar tutoring activities as Week 2. In the Math Circles, children compared a card presenting a set of erasers in a Math Circle to a card presenting a symbolic number (Arabic numeral). In the Math War and Comparing speed, quantities on the cards were in nonsymbolic or symbolic formats. For symbolic format, the child drew a number of dots that corresponds to the symbolic number on the card. Lastly, children completed a review worksheet for Week 3, in which they determined a larger quantity between nonsymbolic and symbolic numbers.

*Week 4.* In Week 4, children practiced comparison between symbolic numbers through similar tutoring activities as Weeks 2 and 3. Children were introduced to an adapted version of Beat Your Score (11) wherein they placed four decks of cards in numerical order for three times (the tutor shuffled the cards for each trial) with an aim of "beating" the time taken for the previous trial. Quantities on each deck of cards were in nonsymbolic (a dot array), mixed (a dot array and numerals), or symbolic formats. In Math War and Comparing speed, quantities on the cards were in symbolic format. Lastly, children completed a review worksheet for Week 4, in which they determined which of two symbolic numbers is larger.

Additional information on the tutoring protocol can be found in Chang et al. (2022).

**Correlations between changes in between-format dissimilarity, changes in format-specific task performance, and fluency gains in children with MD.**
To further examine whether behavioral mapping between nonsymbolic and symbolic number formats may contribute to transfer of learning to arithmetic fluency following INS tutoring, as observed in children with MD, we assessed correlation between changes in between-format dissimilarity (difference in task performance between two number formats) and gains in arithmetic fluency in these children. We additionally tested whether change in arithmetic fluency is related to improved format-specific task performance in each specific format (nonsymbolic or symbolic) in children with MD. Efficiency score was used to assess task performance

### Results

#### No significant effect of head motion

To confirm that any observed differences in task (nonsymbolic, symbolic), time (pre-, post-tutoring), or group (MD, TD) were not confounded by differences in head movement, we performed paired or two-sample *t*-tests on movement parameters. There were no significant differences in movement parameters between nonsymbolic and symbolic number comparison tasks before (*ps* > 0.282) and after (*ps* > 0.247) tutoring, and no significant differences in these parameters between pre- and post-tutoring in nonsymbolic (*ps* > 0.237) and symbolic (*ps* > 0.073) number comparison tasks. Furthermore, there were no significant differences in movement parameters between TD and MD groups for either task or either time point (nonsymbolic at pre-tutoring: *ps* > 0.394; nonsymbolic at post-tutoring: *ps* > 0.307; symbolic at pre-tutoring: *ps* > 0.537; symbolic at post-tutoring: *ps* > 0.597).

**Effects of tutoring on cross-format similarity in numerical processing assessed by accuracy and reaction time on number comparison tasks.**

We examined whether our INS tutoring remediated weak cross-format similarity in numerical processing in children with MD to the level of TD children. As a measure of cross-format similarity in numerical processing, a metric of *between-format dissimilarity* was assessed by absolute difference in behavioral performance between nonsymbolic and symbolic number comparison tasks (|Nonsymbolic – Symbolic|). In addition to efficiency described in the main manuscript, we assessed between-format dissimilarity based on measures of accuracy and reaction time.

For between-format dissimilarity assessed by accuracy, two sample *t*-tests revealed no significant difference between the MD and TD groups at pre-tutoring (*p* = 0.438, Cohen's *d* = 0.22) and between the MD group at post-tutoring and the TD group at pre-tutoring (*p* = 0.915, Cohen's *d* = 0.03) (**Fig S1A**). A post-hoc paired *t*-test also showed no significant tutoring-induced changes in between-format dissimilarity in the MD group (*p* = 0.237, Cohen's *d* = 0.24).

For between-format dissimilarity assessed by reaction time, we found similar patterns of results as that assessed by efficiency. Two sample *t*-tests revealed significant difference between the MD and TD groups at pre-tutoring with moderate effect size (*t*(31.48) = 2.45, *p* = 0.020, Cohen's *d* = 0.70), and no significant difference between the MD group at post-tutoring and the TD group at pre-tutoring (*p* = 0.669, Cohen's *d* = -0.12) (**Fig S1B**). A post-hoc paired *t*-test confirmed significant tutoring-induced changes in between-format dissimilarity in the MD group (*t*(24) = 2.64, *p* = 0.014, Cohen's *d* = 0.53). These results indicate that observed normalization in behavioral between-format dissimilarity assessed by efficiency across children with and without MD were potentially driven by changes in between-format dissimilarity in reaction time, suggesting that INS tutoring may have induced similar processing speed and efficiency between nonsymbolic and symbolic number comparison tasks in children with MD to levels of their TD peers.

Next, we conducted a mixed-design ANOVA with Group (MD, TD) as a between-subject factor and Time (pre-tutoring, post-tutoring) as an within-subject factor to examine whether INS tutoring induced similar or distinct patterns of changes in between-format dissimilarity between the two groups of children. We found a significant Group by Time interaction effect for between-format dissimilarity assessed by reaction time (*F*(1,49) = 8.60, *p* = 0.005, $\eta^2$ = 0.063). Follow-up paired *t*-tests revealed reduced between-format dissimilarity in the MD group (*t*(24) = -2.64, *p* = 0.014, Cohen's *d* = 0.53) and no significant changes in the TD group after tutoring (*p* = 0.242, Cohen's *d* = -0.24). There were no significant main effects of Group or Time for between-format dissimilarity assessed by reaction time (*ps* > 0.064). No significant main effect or interaction was observed for between-format dissimilarity assessed by accuracy (*ps* > 0.227). These results indicate distinct tutoring-induced changes in between-format dissimilarity in reaction time between the MD and TD groups and converge with the patterns of behavioral normalization in children with MD.

**Increased in cross-format similarity in numerical processing was related with gains in arithmetic fluency in children with MD**
We found that tutoring-induced gains in arithmetic fluency in children with MD was negatively correlated with changes in between-format dissimilarity ($r(25) = -0.404$, $p = 0.045$), but not with changes in format-specific task performance ($|rs| < 0.190$, $ps > 0.363$) in these children. These findings indicate that greater reduction in dissimilarity (or increased similarity) between nonsymbolic and symbolic numbers, rather than format-specific performance improvements, may have contributed to transfer of learning to arithmetic fluency in children with MD.

**Effects of tutoring on format-specific number comparison task performance**
We examined whether our INS tutoring remediated number comparison ability in each nonsymbolic and symbolic format in children with MD. We assessed behavioral performance using efficiency, accuracy, and reaction time as described below.

*Efficiency*. Two sample *t*-tests revealed no significant difference between children with MD and TD children at pre-tutoring (nonsymbolic: $p = 0.929$, Cohen's $d = 0.03$; symbolic: $p = 0.102$, Cohen's $d = 0.47$) and between children with MD at post-tutoring and TD children at pre-tutoring (nonsymbolic: $p = 0.139$, Cohen's $d = 0.42$; symbolic: $p = 0.488$, Cohen's $d = 0.20$). These findings suggest that children with MD did not show specific impairments in number comparison ability in a specific format before tutoring.

To further examine whether INS tutoring induced similar or distinct patterns of changes in number comparison ability between the MD and the TD groups, we conducted a mixed-design ANOVA with Group (MD, TD) and Time (pre-tutoring, post-tutoring) factors on efficiency for each number format. In the symbolic comparison task, we found a significant main effect of Time ($F(1,49) = 36.74$, $p < 0.0001$, $\eta^2 = 0.110$) (**Fig S2A**) and marginally significant group effect ($F(1,49) = 3.832$, $p = 0.056$, $\eta^2 = 0.061$) with medium effect size, but interaction was not significant ($p = 0.974$). Similarly, in the nonsymbolic comparison task, we found a significant main effect of Time ($F(1,51) = 17.97$, $p < 0.001$, $\eta^2 = 0.073$), but no other main effect or interaction was significant ($p > 0.335$). Post-hoc paired *t*-tests confirmed that INS tutoring induced significant improvements in efficiency in both groups (MD: nonsymbolic: $t(25) = 2.16$, $p = 0.040$, Cohen's $d = 0.39$; symbolic: $t(25) = 3.81$, $p < 0.001$, Cohen's $d = 0.64$; TD: nonsymbolic: $t(25) = 3.94$, $p < 0.001$, Cohen's $d = 0.75$; symbolic: $t(25) = 4.96$, $p < 0.001$, Cohen's $d = 0.75$). These results suggest that INS tutoring induced improvements on both symbolic and nonsymbolic comparison ability across the MD group as the TD group.

*Accuracy*. Planned paired t-test did not show any significant improvements in the MD group for both tasks (nonsymbolic: $p = 0.083$, Cohen's $d = -0.37$; symbolic: $p = 0.248$, Cohen's $d = 0.22$) (**Fig S2B**). Also, two-sample t-tests revealed no significant differences between the MD and TD groups at pre-tutoring (nonsymbolic: $p = 0.323$, Cohen's $d = -0.27$; symbolic: $p = 0.141$, Cohen's $d = 0.42$) and between the MD group at post-tutoring and the TD group at pre-tutoring (nonsymbolic: $p = 0.988$, Cohen's $d = 0.004$; symbolic: $p = 0.451$, Cohen's $d = 0.21$).
To further unpack similar learning trajectories between the MD and the TD groups found with efficiency measure in the main text, we conducted the same mixed-design ANOVA, with Group (MD, TD) as the between-subject factor and Time (Pre, Post) as the within-subject factor for each comparison task with accuracy. In both nonsymbolic and symbolic number comparison tasks, we did not find any significant main effect or interaction (nonsymbolic: $ps > 0.178$; symbolic: $ps > 0.262$). Post-hoc *t*-test confirmed that both MD and TD groups showed no significant changes in accuracy (nonsymbolic: $ps > 0.082$, Cohen's $|d| < 0.37$; symbolic: $ps > 0.248$, Cohen's $|d| < 0.23$).

*Reaction time*. With reaction times, planned paired t-tests showed significant reduction in reaction times in the MD group for both tasks (nonsymbolic: $t(25) = -2.45$, $p = 0.022$, Cohen's $d = -0.40$; symbolic: $t(24) = -3.78$, $p < 0.001$, Cohen's $d = -0.55$) (**Fig S2C**). However, as accuracy and efficiency measures, two-sample t-test with reaction times revealed no significant group differences at pre-tutoring (nonsymbolic: $p = 0.429$, Cohen's $d = -0.22$; symbolic: $p = 0.132$, Cohen's $d = -0.43$) and between the MD group at post-tutoring and the TD group at pre-tutoring (nonsymbolic: $p = 0.409$, Cohen's $d = 0.23$; symbolic: $p = 0.604$, Cohen's $d = 0.15$). These results showed that tutoring-induced changes in efficiency measure was mainly driven by changes in reaction times in the MD groups and confirmed the results that the MD group did not show any deficits in performing number comparison tasks.
To further unpack similar learning trajectories between the MD and the TD groups found with efficiency measure in the main text, we conducted the same mixed-design ANOVA, with Group (MD, TD) as the between-subject factor and Time (Pre, Post) as the within-subject factor for each comparison task with reaction time. Here we found significant time effects with reaction times for nonsymbolic number comparison task ($F(1,51) = 18.34$, $p < .001$, $\eta^2 = 0.059$). No other main effect or interaction was significant ($ps > 0.290$). For symbolic number comparison task, we found significant time effects with reaction times ($F(1,49) = 22.12$, $p < 0.001$, $\eta^2 = 0.067$). No other effects were significant ($ps > 0.136$). Post-hoc *t*-tests revealed that both groups showed significant reduction in reaction times for both tasks (TD: nonsymbolic: $t(26) = -3.94$, $p < 0.001$, Cohen's $d = -0.64$; symbolic: $t(25) = -2.83$, $p = 0.009$, Cohen's $d = -0.51$; MD: nonsymbolic: $t(25) = -2.45$, $p = 0.022$, Cohen's $d = -0.40$; symbolic: $t(24) = -3.78$, $p < 0.001$, Cohen's $d = -0.55$). These results confirmed the results with efficiency in the main text that INS tutoring leads to similar levels of improvement across the MD and TD groups.

**INS tutoring leads to distinctive pattern of changes in arithmetic fluency in children with and without MD**
To further examine whether INS tutoring induced similar or distinct patterns of changes in arithmetic fluency between MD and TD groups, we conducted a mixed-design ANOVA with Group (MD, TD) and Time (pre-tutoring, post-tutoring) factors. We found significant main effect of Group ($F(1,51) = 43.51$, $p < 0.001$, $\eta^2 = 0.39$) and interaction effect between Group and Time ($F(1,51) = 7.537$, $p < 0.001$, $\eta^2 = 0.04$). Main effect of Time was not significant ($p = .121$). Post-hoc paired *t*-tests revealed significant increases in arithmetic fluency in the MD group ($t(25) = 3.41$, $p = 0.002$, Cohen's $d = 0.90$) and no significant changes in arithmetic fluency in the TD group ($p = 0.453$, Cohen's $d = 0.13$). Our results suggest that INS tutoring focused on integration of nonsymbolic and symbolic number representations induced distinct patterns of changes in arithmetic fluency between children with MD and TD children, with significant improvements observed in children with MD.

**Similar learning trajectories between the MD and TD groups: Whole-brain ANOVA analysis**
We examined whether INS tutoring induces similar or different patterns of plasticity in cross-format NRS in children with MD and TD children. We performed a whole-brain mixed ANOVA with Group (MD, TD) as a between-subjects factor and Time (Pre, Post) as a within-subjects factor to assess the effect of tutoring that is similar or different between groups. Main effects of Group and Time are described below (see main manuscript for results of interaction effect). Follow-up ROI analyses were performed to further identify the directionality of observed effects.

***Group effect.*** We observed a significant main effect of Group in the bilateral intraparietal sulcus (IPS; 42, -44, 46; -36, -50, 52) and left hippocampus (Hipp; -28, -16, -12), with higher cross-format NRS in the TD compared to the MD group (**Table S7**). These findings of group effect are consistent with the whole brain analysis contrasting TD and MD groups at pre-tutoring (**S3** Table).

***Time effect.*** We observed a significant main effect of Time (i.e., effect of tutoring) in different brain regions, with cross-format NRS after tutoring increased especially in the left Hipp (-16, -20, 2) and decreased in the right frontal pole (28, 54, 8) (**Fig S3A**, **Table S8**). Follow-up paired *t*-tests confirmed that the MD showed significant tutoring-related changes in cross-format NRS in the left Hipp (MD group: $t(18) = 2.94$, *FDR-corrected* $p = 0.014$, Cohen's $d = 0.86$) and right frontal pole (MD group: $t(18) = -2.90$ , $p = 0.014$, Cohen's $d = -0.72$), but not in the TD group (left Hipp: $t(20) = 2.26$, *FDR-corrected* $p = 0.821$, Cohen's $d = 0.70$; right frontal pole: $t(20) = -2.35$, *FDR-corrected* $p =.0821$, Cohen's $d = -0.70$). An additional whole-brain paired *t*-test contrasting pre- and post-tutoring across all children showed similar patterns of results (**S**9 Table, **S**4A Fig). Together, these results demonstrate that INS tutoring induce significant changes in cross-format NRS in hippocampal and frontal cortical regions in the MD group.

***Broader involvement of the hippocampus in children with MD compared to TD children***. It is worth mentioning that multiple hippocampal regions identified in the analysis of Time and Group effects from whole-brain ANOVA and the analysis of pre-tutoring group difference in whole-brain two-sample t-test (described in the main manuscript) were anatomically distinct (**S**5 Fig). Specifically, the region identified from the t-test was located in the lateral-posterior side of the hippocampus, the region identified from the Group effect was in the middle of the hippocampus, and lastly, the region identified from the Time effect was located in the posterior side of the hippocampus. The MD group showed increased cross-format NRS or remediation of cross-format NRS in all middle and posterior regions of the hippocampus after tutoring, whereas the TD group showed increased cross-format NRS only in the posterior hippocampus. These findings indicate children with MD show tutoring-induced changes in broader regions of hippocampus compared to TD children. Considering the key role of the hippocampus in integration of related memory (Eichenbaum, 2004; Giovanello et al., 2004; Olsen et al., 2012; Ranganath, 2010; Staresina & Davachi, 2009), these findings suggest that INS tutoring contributed to enhanced binding of neural representations across nonsymbolic and symbolic numbers, especially in children with low math ability. Combined with our behavioral findings of normalization of cross-format similarity in children with MD, our results may further suggest a role of hippocampus in enhancing learning of the relations of two number formats (Chang et al., 2022) in these children.

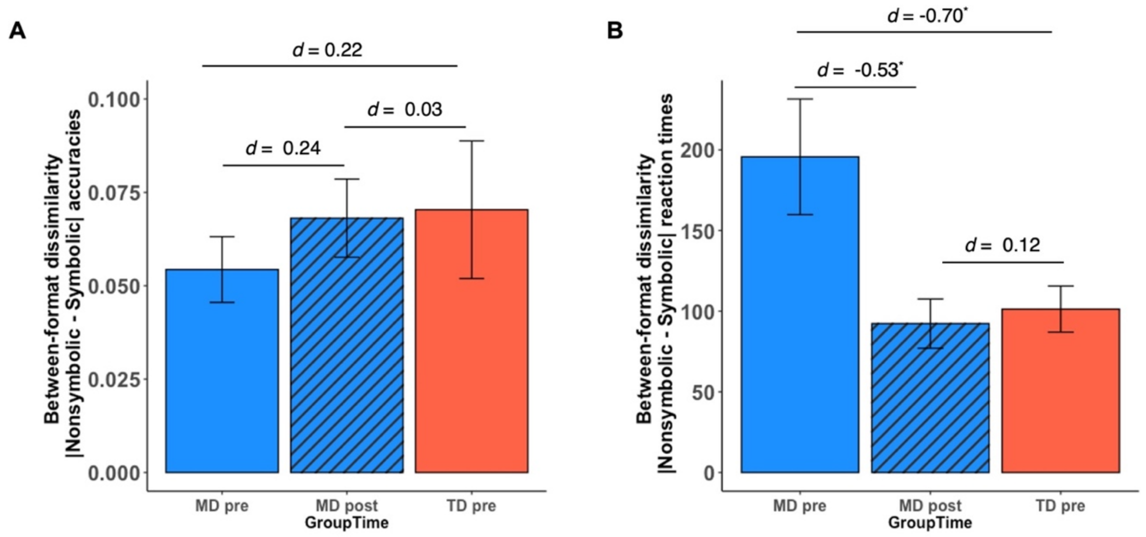

**Fig. S1. INS tutoring normalized weak cross-format similarity in behavioral performance between nonsymbolic and symbolic number formats.**

Between-format dissimilarity was measured by the absolute difference in behavioral performance between nonsymbolic and symbolic number comparison tasks (|Nonsymbolic – Symbolic|). Higher scores represented higher between-format dissimilarity (or lower cross-format similarity) in behavioral performance. (A) Between-format dissimilarity measured by accuracy. Effect sizes indicate that group difference in between-format similarity at pre-tutoring ($d$ = 0.22) was greater than difference in between-format similarity between the MD group at post-tutoring and the TD group at pre-tutoring ($d$ = 0.03). (B) Between-format dissimilarity measured by reaction time. Between-format dissimilarity was significantly higher in the MD, compared to the TD, group at pre-tutoring ($p$ = 0.020, Cohen's $d$ = -0.70). INS tutoring reduced between-format dissimilarity in the MD group at post-tutoring to the level of the TD group at pre-tutoring ($p$ = 0.669, Cohen's $d$ = 0.12). $^{*}p < 0.05$, $d$ = Cohen's $d$.

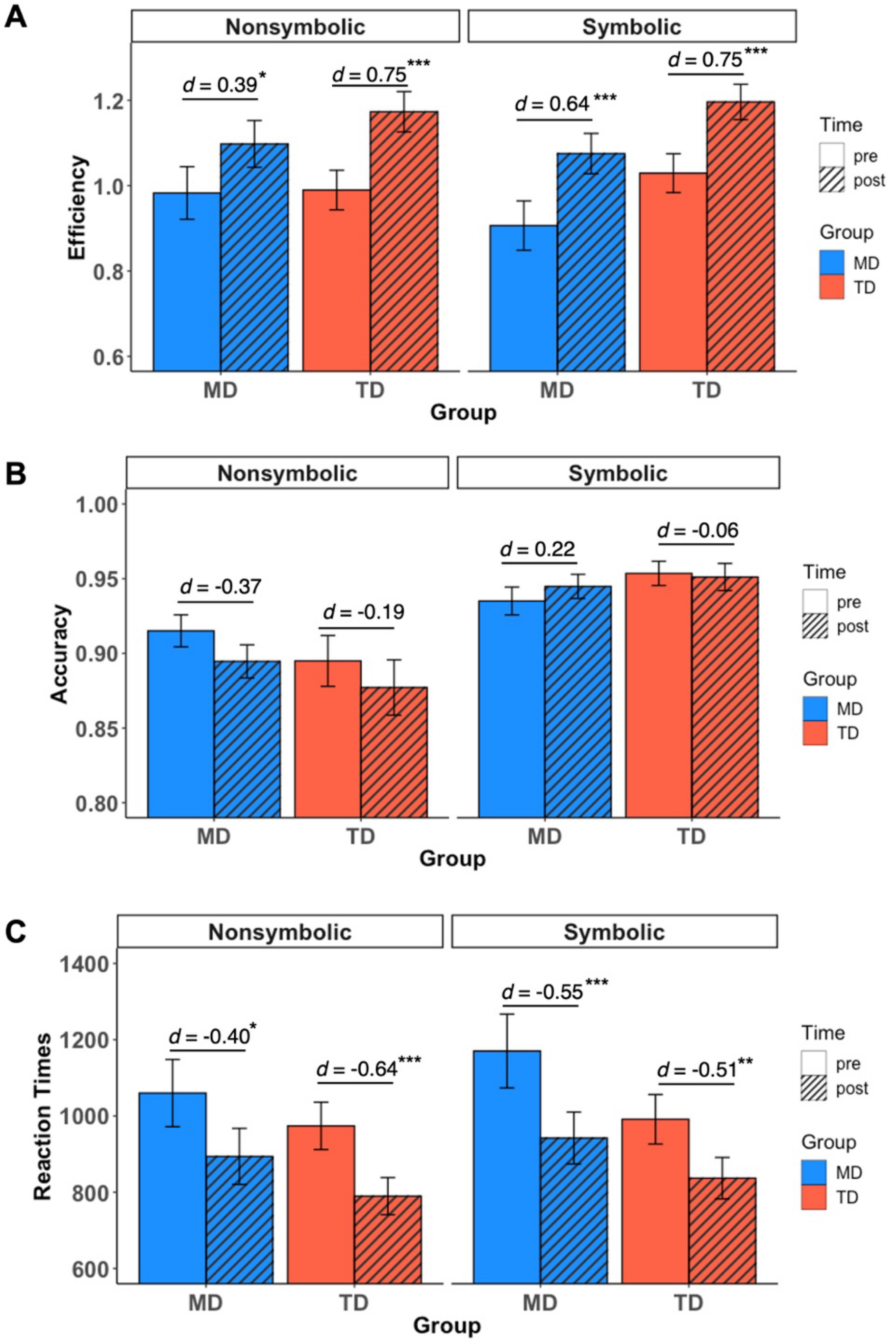

**Fig. S2. Tutoring induced changes in number comparison task performance in children with MD and TD children.**

(A) INS tutoring significantly increased efficiency scores in nonsymbolic and symbolic comparison tasks in both MD (blue bars) and TD (red bars) groups. Efficiency scores were obtained by dividing accuracy by median reaction time. (B) INS tutoring did not significantly change accuracy on nonsymbolic and symbolic comparison tasks in both MD and TD groups. C. INS tutoring significantly decreased median reaction times for nonsymbolic and symbolic comparison tasks in both MD and TD groups. $^{*}p < 0.05$, $^{**}p < 0.01$, $^{***}p < 0.001$, $d$ = Cohen's *d.*

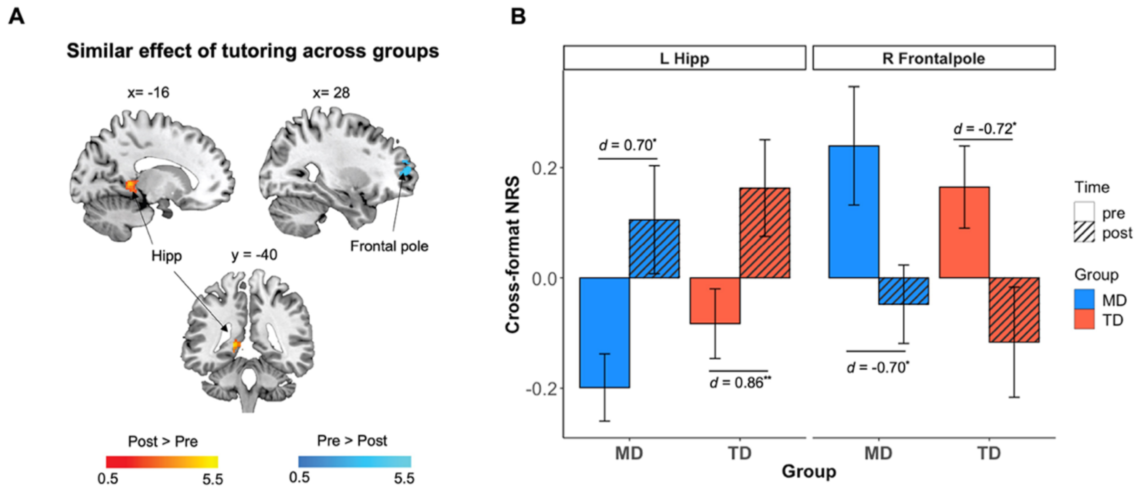

**Fig. S3. Tutoring induced similar patterns of changes in cross-format NRS in children with MD and TD children.**

(**A**) A whole-brain mixed ANOVA with a within-subject factor Time (pre-tutoring, post-tutoring) and a between-subject factor Group (MD, TD) revealed a significant main effect of Time in multiple brain regions including increased cross-format NRS in the left hippocampus (Hipp) and decreased cross-format NRS in the right frontal pole following tutoring. (**B**) Follow-up regional analyses confirmed significant changes in the left Hipp and right frontal pole in MD (blue bars) but not in TD (red bars) groups. **FDR-corrected* $p < 0.05$. $d$ = Cohen's $d$. L, Left; R, Right.

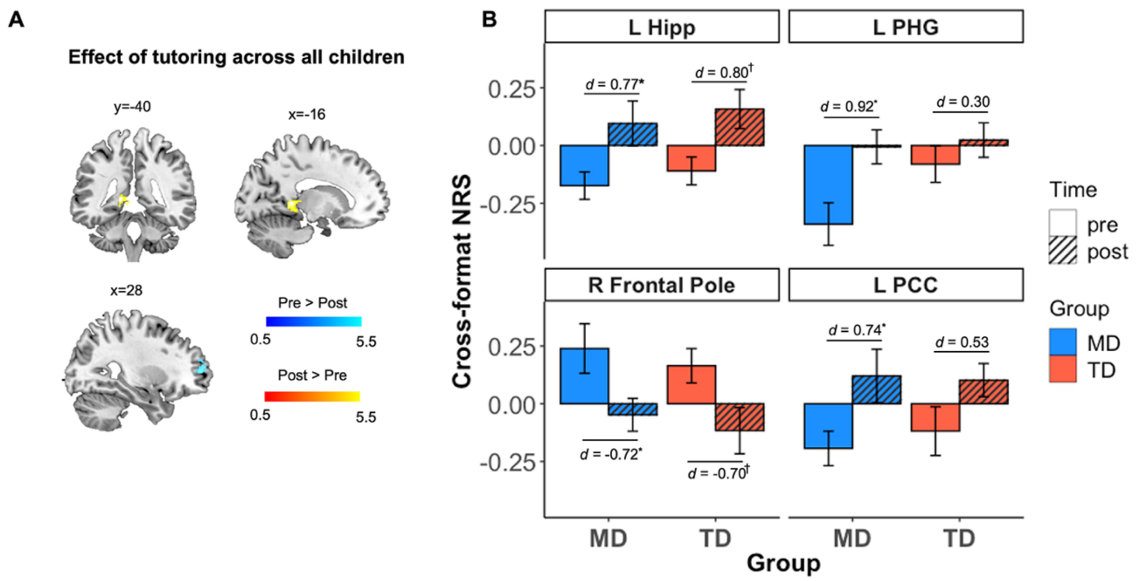

**Fig. S4. Tutoring induced changes in cross-format NRS in the left hippocampus and right frontal pole across all children.**

(**A**) A whole-brain paired *t*-test contrasting pre- and post-tutoring across all children revealed significantly increased cross-format NRS in the left hippocampus (Hipp), left parahippocampal gyrus (PHG), and posterior cingulate gyrus (PCC) and significantly decreased cross-format NRS in the right frontal pole after tutoring. (**B)** Follow-up regional *t*-tests confirmed a significant increase in cross-format NRS in the left Hipp and significant decrease in cross-format NRS in the right frontal pole for MD (blue bars) and TD (red bars) groups. Significant increases in cross-format NRS in the left PHG and left PCC were observed in children with MD but not in TD children. All p-values are *FDR-corrected;* **p* < .05, ***p* < .01, *d* = Cohen's *d.* L, Left; R, Right.

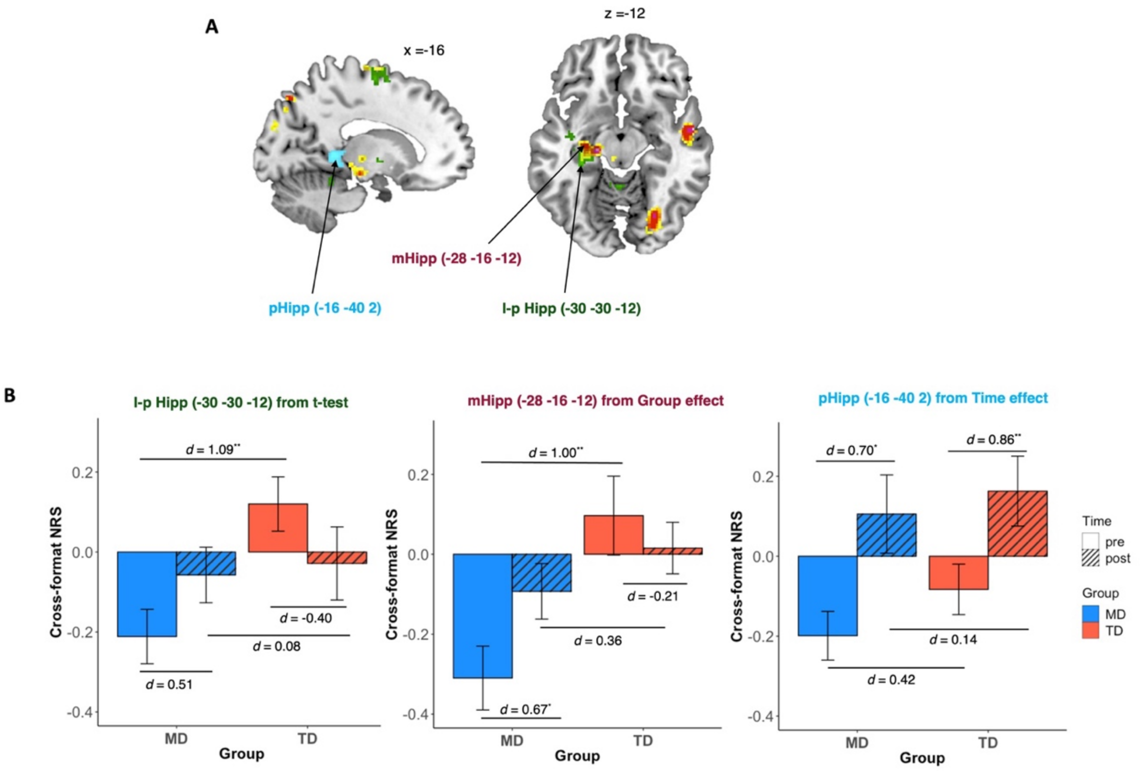

**Fig. S5. Tutoring induced distinct patterns of changes in cross-format NRS in subdivisions of the left hippocampus in children with MD and TD children.**

(**A**) The left hippocampus regions identified from whole-brain paired *t*-test and group (MD, TD) by time (pre, post) ANOVA were anatomically distinct. The left hippocampus region from *t*-test contrasting pre- and post-tutoring was located in the lateral-posterior side of the hippocampus (l-pHipp, green), the region from the main effect of group from ANOVA was located in the middle hippocampus (mHipp, red) and the region from the main effect of time from ANOVA was located in the posterior hippocampus (pHipp, blue). (**B**) Follow-up regional analyses clarified that l-pHipp, mHipp and pHipp showed distinct patterns of change in cross-format NRS following tutoring across MD and TD groups. While children with MD showed significant increases in cross-format NRS in all regions, TD children showed significant increases in cross-format NRS only in the pHipp. All p-values are *FDR-corrected;* *$p < 0.05$, **$p < 0.01$, $d$ = Cohen's $d$

**Table S1.** Summary of previous studies of number sense intervention in children with or without mathematical disabilities

| | Age | | Sample Size | | | Training types | | | Intervention duration | Imaging |
|---|---|---|---|---|---|---|---|---|---|---|
| Study | M (yr.) | SD (mo.) | TD | TD Cntrl | MD | NonSym | Sym + Nonsym | Sym | | |
| Wilson et al (2006) (Wilson et al., 2006) | 7-10 yrs | | | | 13 | O | O | | 5-weeks, 4 days/week, 30 min/session | n/a |
| Opfer & Siegler (2007)(Opfer & Siegler, 2007) | 8.20 | 7.20 | 61 | | | | O | | 3 trial blocks consisted of 10 items per each | n/a |
| Booth & Siegler (2008) (Booth & Siegler, 2008) | 7.20 | 4.80 | 78 | 27 | | | O | | 1-week, 3 sessions, 10-15/session | n/a |
| Siegler & Ramani (2008) (Siegler & Ramani, 2008) | 4.60 | 3.60 | 36 | | | | O | | 2-weeks, 4 sessions, 15min/session | n/a |
| | 4.70 | 5.04 | | 18 | | | | | | |
| Ramani & Siegler (2008) (Ramani & Siegler, 2008) | 4.75 | 5.52 | 68[1] | | | | O | | 2-weeks, 4 sessions, 15-20min/session | n/a |
| | 4.75 | 4.92 | | 56 | | | | | | |
| Siegler & Ramani (2009) (Siegler & Ramani, 2009) | 4.66 | 5.40-5.52 | 59[1] | | | | O | | 3-weeks, 5 sessions, 15-20min/session | n/a |
| | 4.66 | 6.24 | 29[1] | | | | | O | | |
| Wilson et al. (2009) (Wilson et al., 2009) | 5.60 | 4.80 | 53[1] | | | O | O | | 6 sessions, 20 min/session | n/a |
| Ramani & Siegler (2011) (Ramani & Siegler, 2011) | 4.00 | 3.84-4.56 | 59 | | | | O | | 3-weeks, 5 sessions, 15-20min/session | n/a |
| | 4.00 | 4.80 | 29 | | | | | O | | |
| **Kucian et al (2011) (Kucian et al., 2011)** | **9.50** | **13.20** | | | **16** | O | O | | **5-weeks, 5 sessions/week, 15 min** | **fMRI** |
| | **9.60** | **9.60** | **16** | | | | | | | |
| Ramini et al. (2012) (Ramani et al., 2012) | 4.58 | 6.12 | 34[1] | | | | O | | 3-4 weeks, 6 sessions, 20-25 min/session | n/a |
| | 4.17 | 6.84 | | 28[1] | | | | | | |
| Obersteiner et al. (2013) (Obersteiner et al., 2013) | 6.91 | 4.68 | 35 | | | O | O | | 4-weeks, 30 min,10 sessions | n/a |
| | | | 39 | | | O | O | O | | |
| | | | 39 | | | | O | O | | |
| | | | | 34 | | | | | | |

[1] Low SES children

| | | | | | | | | | | |
|---|---|---|---|---|---|---|---|---|---|---|
| Hyde et al. (2014) (Hyde et al., 2014) | 6.89 | 2.59 | 96 | | | O | | | 2 sets of practice (60 problems) | n/a |
| Kuhn & Hollings (2014) (Kuhn & Holling, 2014) | 9.00 | 8.40 | 20 | 20 | | O | | O | 3-weeks, 15 sessions, 20 min/session | n/a |
| Honore & Noel (2016) (Honore & Noel, 2016) | 5.75 | 3.79 | 19 | 18 | | O | | | 6-weeks, 10 sessions, 30 min/session | n/a |
| | | | 19 | | | | O | | | |
| Sella et al (2016) (Sella et al., 2016) | 5.17 | 8.00 | 23 | | | O | O | | 10-weeks, 2 session/week, 20min/session | n/a |
| | 5.00 | 7.00 | | 22 | | | | | | |
| Elofsson et al. (2016) (Elofsson et al., 2016) | 5.37 | 3.72 | 54 | 60 | | | O | | 3 weeks, 6 sessions, 10 min/session | n/a |
| Maertens et al. (2016) (Maertens et al., 2016) | 5.44 | 3.48 | 47 | 63 | | O | | O | 3 weeks, 6 sessions, 10 min/session | n/a |
| | 5.32 | 3.72 | 41 | | | | O | | | |
| Park et al.(2016) (Park et al., 2016) | 4.87 | 4.8 | 51 | 52 | | O | | | 2-3 weeks, 10 sessions, 12min/session | n/a |
| Van Herwegen et al. (2017) (Van Herwegen et al., 2017) | 3.77 | 7.20 | 20 | | | O | O | | 5-weeks, 10min/day | n/a |
| | 3.82 | 6.06 | | 18 | | | | | | |
| **Looi et al. (2017) (Looi et al., 2017)** | **9.48** | **7.3** | | | **6** | | tRNS | | **5 weeks, 2 sessions/week, 20 min/session** | **tRNS** |
| | | | | | **6** | | Sham | | | |
| Ramani et al. (2017) (Ramani et al., 2017) | 6.01 | 4.31 | 27 | 27 | | | O | | 10 sessions, 10-15min/session | n/a |
| **Michels et al. (2018) (Michels et al., 2018)** | **9.50** | **8.40** | | | **15** | O | O | | **5-weeks, 5 sessions/week, 15 min** | **fMRI** |
| | **9.50** | **9.60** | **16** | | | | | | | |
| Kim et al. (2018) (Kim et al., 2018) | 7.70 | 3.60 | 22 | | 24 | O | O | | 6-weeks, 30 sessions, 24 min/session | n/a |
| Szkudlarek & Brannon (2018) (Szkudlarek et al., 2021) | 4.57 | 7.32 | 53 | | | O | | | 10 sessions, 12 min/session | n/a |
| | 4.37 | 7.44 | | | [2]27 | | | | | |
| | 4.61 | 6.24 | 52 | | | | | O | | |
| | 4.48 | 6.12 | | | 29 | | | | | |
| | 4.58 | 6.72 | | 52 | | | | | | |
| | 4.41 | 7.08 | | | 31 | | | | | |
| | 3.62 | 3.99 | | | 19 | O | | | 5-weeks, 10min/day | |

[2] Among TD children, post-tested a part of children with relatively low math

| | | | | | | | | | | |
|---|---|---|---|---|---|---|---|---|---|---|
| Van Herwegen et al. (2018) (Van Herwegen et al., 2018) | 3.68 | 4.52 | | | 19 | | | O | | |
| | 3.77 | 3.91 | 20 | | | | | | | |
| Whyte & Bull (2018) (Whyte & Bull, 2008) | 3.80 | 4.00 | 32 | 13 | | | O | | 4 sessions, 25 min/session | n/a |
| Ramani et al. (2020) (Ramani et al., 2020) | 6.02 | 3.94 | 47 | | | | O | | 10 sessions, 10-15 min/session | n/a |
| | 5.94 | 3.39 | | 53 | | | | | | |
| Libertus et al. (2020) (Libertus et al., 2020) | 6.17 | 8.1 | 33 | 35 | | O | | | 5 weeks, 16 sessions, 15min/ session | n/a |
| Vanbecelaere et al. (2020) (Vanbecelaere et al., 2020) | 6.37 | 5.04 | 109 | 223 | | O | O | | 6-weeks, 6 sessions, 50min/session | n/a |
| Bugden et al. (2021) (Bugden et al., 2021) | 9.75 | 7.44 | 53 | | | O | | | 6 days, 20-30min/session | n/a |
| | 9.73 | 8.64 | | 58 | | | | | | |
| Tobia et al. (2021) (Tobia et al., 2021) | 4.76 | 3.24 | | | 36 | O | | | 7-weeks, 3 sessions/week, 45 min/session | n/a |
| | | | | | 29 | | O | O | | |
| | | | | | 24 | | | | | |
| | 4.77 | 3.6 | 27 | | | O | | | | |
| | | | 23 | | | | O | O | | |

Abbreviations: M, mean; SD; standard deviation; TD, typically-developing; cntl, controls; MD, mathematical disabilities or children with low math ability; Nonsym, nonsymbolic; Sym, symbolic.

**Table S2.** Demographics and standardized assessment scores in the MD and TD grous.

| | | MD | | | TD | | | MD vs. TD |
|---|---|---|---|---|---|---|---|---|
| | | *M* | *SD* | range | *M* | *SD* | range | **p*-values |
| **Age** | | 8.33 | 0.67 | 7.40-10.01 | 8.04 | 0.46 | 7.17-9.09 | 0.130 |
| **Gender** | | 9 Males, 10 Females | | | 9 Males, 12 Females | | | |
| **WASI** | FSIQ | 103.40 | 11.4 | 88-124 | 106.70 | 12.6 | 86-133 | 0.399 |
| | VIQ | 105.68 | 11.5 | 83-126 | 107.80 | 12.6 | 89-133 | 0.588 |
| | PIQ | 101.21 | 15.4 | 84-140 | 104.57 | 14.9 | 83-131 | 0.488 |
| **WJ-III (math)** | Math fluency | 86.00 | 3.3 | 80-90 | 102.43 | 8.8 | 92-121 | <0.001 |
| **WJ-III (reading)** | Letter word Identification | 108.37 | 11.1 | 95-130 | 109.71 | 7.1 | 93-125 | 0.656 |
| | Word attack | 104.16 | 8.8 | 90-121 | 108.00 | 4.0 | 101-115 | 0.094 |
| **AWMA (working memory)** | Digit recall | 96.79 | 14.9 | 70-129 | 99.38 | 12.0 | 76-125 | 0.552 |
| | Word recall | 87.11 | 16.3 | 64-115 | 90.62 | 16.9 | 64-115 | 0.507 |
| | Backward digit recall | 99.16 | 15.7 | 81-143 | 102.38 | 11.8 | 81-122 | 0.473 |
| | Block recall | 84.32 | 14.0 | 67-109 | 91.43 | 12.4 | 74-111 | 0.099 |
| | Spatial recall | 99.16 | 21.4 | 64-128 | 106.60 | 14.8 | 83-125 | 0.216 |

*Note, *p* values from two sample t-test

**Table S3.** Group differences between the MD and TD groups in cross-format NRS at pre-tutoring.

| Region | Cluster Size | T-value | MNI coordinates x | y | z |
|---|---|---|---|---|---|
| **Pre-tutoring** | | | | | |
| ***TD pre > MD pre*** | | | | | |
| L SPL/IPS | 286 | 5.11 | -36 | -50 | 52 |
| L SPL/IPS | | 4.14 | -26 | -54 | 54 |
| L SPL | | 3.79 | -30 | -54 | 62 |
| R postCG | 139 | 4.78 | 54 | -22 | 46 |
| R SMG | | 4.09 | 48 | -32 | 50 |
| R postCG | | 3.06 | 62 | -18 | 42 |
| R Cerebellum[3] | 120 | 4.64 | 22 | -76 | -24 |
| R FG | | 4.03 | 34 | -70 | -22 |
| R Cerebellum[1] | | 3.46 | 12 | -76 | -24 |
| R COC | 127 | 4.46 | 44 | -12 | 16 |
| R COC | | 3.42 | 50 | -16 | 20 |
| L Cerebellum[1] | 111 | 4.46 | -2 | -48 | -14 |
| L PHG | | 3.61 | -22 | -38 | -16 |
| L Cerebellum[1] | | 3.45 | -14 | -44 | -18 |
| L MFG/FEF | 363 | 4.27 | -32 | 2 | 64 |
| L SFG | | 4.11 | -14 | 2 | 62 |
| L preCG | | 3.92 | -30 | -10 | 60 |
| L Putamen | 102 | 4.16 | -26 | -2 | 14 |
| L Insular | | 3.02 | -30 | 6 | 14 |
| R preCG | 164 | 4.15 | 2 | -24 | 68 |

[3] Four cerebellar ROIs were excluded from a multivariate classification analysis due to insufficient coverage of the cerebellum during data acquisition.

| | | | | | |
|---|---|---|---|---|---|
| L preCG | | 3.17 | -4 | -24 | 58 |
| L Pallidum | 378 | 3.87 | -22 | -10 | 0 |
| L Hippocampus | | 3.87 | -30 | -30 | -12 |
| L Thalamus | | 3.78 | -20 | -26 | 4 |
| ***MD pre > TD pre*** | | | | | |
| No significant clusters | | | | | |

Abbreviations: COC, Central Opercular Cortex; FEF, Frontal Eye Fields; FG, Fusiform Gyrus; IPS, Intraparietal Sulcus; MFG, Middle Frontal Gyrus; PHG, Parahippocampal gyrus; postCG, Postcentral Gyrus; preCG, Precentral Gyrus; SMG, Supramarginal Gyrus; SFG, Superior Frontal Gyrus; SPL, Superior Parietal Lobule; L, Left; R, Right.

**Table S4.** Two-sample t-test between cross-format NRS in the MD group at pre- and post-tutoring and cross-format NRS in the TD group at pre-tutoring

| | **MD pre vs. TD pre** | | | **MD post vs. TD pre** | | |
|---|---|---|---|---|---|---|
| ROI [MNI coordinates]: | *t* | *p*⁺ | Cohen's *d* | *t* | *p*⁺ | Cohen's *d* |
| L SPL/IPS [-36 -50 52] | -4.56*** | 0.001 | 1.41 | -2.10 | 0.204 | 0.66 |
| L SPL/IPS [-26 -54 54] | -3.86*** | 0.001 | 1.20 | -1.60 | 0.294 | 0.50 |
| L SPL [-30 -54 62] | -3.47** | 0.002 | 1.08 | -2.02 | 0.204 | 0.63 |
| R postCG [52 -22 46] | -4.42*** | 0.001 | 1.42 | -2.09 | 0.204 | 0.68 |
| R SMG [48 -32 50] | -3.76** | 0.002 | 1.20 | -1.68 | 0.290 | 0.54 |
| R postCG [62 -18 42] | -3.15** | 0.004 | 1.01 | -0.69 | 0.554 | 0.22 |
| R FG [34 -70 -22] | -3.08** | 0.005 | 0.98 | -1.19 | 0.405 | 0.38 |
| R COC [44 -12 16] | -4.07*** | 0.001 | 1.28 | -1.26 | 0.389 | 0.40 |
| R COC [50 -16 20] | -2.69* | 0.012 | 0.85 | -0.30 | 0.807 | 0.09 |
| L PHG [-22 -38 -16] | -2.55* | 0.016 | 0.82 | -0.94 | 0.482 | 0.30 |
| L MFG/FEF [-32 2 64] | -3.74** | 0.002 | 1.16 | -2.09 | 0.204 | 0.66 |
| L SFG [-14 2 62] | -3.96*** | 0.001 | 1.26 | -1.50 | 0.322 | 0.48 |
| L preCG [-30 -10 60] | -3.62** | 0.002 | 1.17 | -2.36 | 0.204 | 0.76 |
| L Putamen [-26 -2 14] | -4.25*** | 0.001 | 1.36 | -1.31 | 0.389 | 0.42 |
| L Insula [-30 6 14] | -3.18** | 0.004 | 1.00 | -0.87 | 0.488 | 0.27 |
| R preCG [2 -24 68] | -3.86*** | 0.001 | 1.20 | -0.97 | 0.482 | 0.31 |
| L PreCG [-4 -24 58] | -2.57* | 0.015 | 0.80 | -0.16 | 0.871 | 0.05 |
| L Pallidum [-22 -10 0] | -3.46** | 0.002 | 1.10 | -0.92 | 0.482 | 0.29 |
| L Hippocampus [-30 -30 -12] | -3.45** | 0.002 | 1.09 | -1.83 | 0.251 | 0.58 |
| L Thalamus [-20 -26 4] | -3.08** | 0.005 | 0.98 | -0.68 | 0.554 | 0.22 |

*Notes*: ROIs were defined from a whole-brain analysis comparing cross-format NRS between the MD and TD groups at pre-tutoring (Table S3).*$p < .05$, **$p < .01$, ***$p < .001$. ⁺All *p*-values are corrected for multiple comparison using FDR correction. Cohen's *d* presents absolute value.

**Table S5.** Brain regions showing significant interaction between group (MD vs. TD) and time (pre- vs. post- tutoring) on cross-format NRS in Group X Time ANOVA.

| | | | MNI coordinates | | |
|---|---|---|---|---|---|
| **Region** | **Cluster Size** | **T-value** | **x** | **y** | **z** |
| ***MD (post>pre) > TD (post>pre)*** | | | | | |
| L Cerebellum[4] | 118 | 5.30 | -24 | -64 | -28 |
| L Cerebellum | | 3.12 | -36 | -60 | -28 |
| L PHG | 112 | 4.25 | -20 | -34 | -14 |
| L Cerebellum | | 4.15 | -14 | -46 | -18 |
| L Cerebellum | | 3.48 | -2 | -48 | -14 |
| L Caudate | 115 | 4.06 | -14 | 24 | 4 |
| L Caudate | | 3.75 | -14 | 18 | 10 |
| L Caudate | | 3.67 | -8 | 18 | 0 |
| L Premotor | 98 | 3.74 | -14 | 2 | 60 |
| L Premotor | | 3.69 | -20 | -10 | 66 |
| L Premotor | | 3.53 | -10 | -12 | 60 |
| ***TD (post>pre) > MD (post>pre)*** | | | | | |
| No significant clusters | | | | | |

Abbreviations: PHG, Parahippocampal Gyrus; L, Left; R, Right.

[4] Cerebellar ROI was excluded a multivariate classification analysis due to insufficient coverage of the cerebellum during data acquisition.

**Table S6.** Paired t-test between cross-format NRS at pre- and post-tutoring in each MD and TD group.

| | MD pre vs. post | | | TD pre vs. post | | |
|---|---|---|---|---|---|---|
| ROI [MNI coordinates]: | *t* | $p^{+}$ | *d* | *t* | $p^{+}$ | *d* |
| L PHG [-20 -34 -14] | -2.42 | 0.037 | -0.79 | 2.03 | 0.097 | 0.59 |
| L Caudate [-14 24 4] | -2.56 | 0.035 | -0.58 | 2.35 | 0.074 | 0.71 |
| L Caudate [-14 18 10] | -3.27 | 0.015 | -0.75 | 1.20 | 0.304 | 0.37 |
| L Caudate [-8 18 0] | -4.16 | 0.004 | -1.15 | 1.16 | 0.304 | 0.35 |
| L Premotor [-14 2 60] | -1.35 | 0.193 | -0.51 | 4.10 | 0.004 | 1.18 |
| L Premotor [-20 -10 66] | -1.91 | 0.083 | -0.69 | 2.31 | 0.074 | 0.65 |
| L Premotor [-10 -12 60] | -2.84 | 0.026 | -0.75 | 0.90 | 0.379 | 0.26 |

**Table S7.** Brain regions showing significant main effect of group (MD vs. TD) on cross-format NRS in Group X Time ANOVA.

| Region | Cluster Size | T-value | MNI coordinates x | y | z |
|---|---|---|---|---|---|
| ***TD > MD*** | | | | | |
| L IPS | 333 | 5.55 | -36 | -50 | 52 |
| L SPL | | 5.00 | -36 | -40 | 58 |
| L SPL | | 5.00 | -36 | -40 | 58 |
| R IPS | 217 | 5.52 | 42 | -44 | 46 |
| R IPL | | 4.89 | 48 | -34 | 50 |
| R SPL | | 3.48 | 36 | -38 | 40 |
| R STG | 212 | 5.41 | 56 | -4 | -12 |
| R MTG | | 4.00 | 50 | -10 | -18 |
| R STG | | 3.61 | 50 | -14 | -4 |
| L Hippocampus | 352 | 5.24 | -20 | -20 | -10 |
| L Hippocampus$^{\dagger}$ | 72 | 4.74 | -20 | -22 | -12 |
| L Hippocampus$^{\dagger}$ | | 4.30 | -28 | -16 | -12 |
| L Hippocampus$^{\dagger}$ | 13 | 4.30 | -22 | -30 | -4 |
| L Hippocampus$^{\dagger}$ | 1 | 3.06 | -16 | -26 | -8 |
| L MFG | 105 | 5.04 | -28 | 4 | 66 |
| L MFG | | 4.39 | -28 | 10 | 60 |
| R Cerebellum[5] | 278 | 5.02 | 22 | -74 | -22 |
| R Cerebellum | | 4.65 | 28 | -80 | -24 |
| R FG | | 4.64 | 26 | -72 | -12 |
| L Putamen | 134 | 4.99 | -28 | -4 | 14 |

[5] Cerebellar ROI was excluded a multivariate classification analysis due to insufficient coverage of the cerebellum during data acquisition.

| | | | | | |
|---|---|---|---|---|---|
| L LOC | 88 | 4.76 | -34 | -84 | 22 |
| L LOC | | 4.27 | -20 | -88 | 26 |
| L Cuneal Cortex | | 3.31 | -10 | -88 | 26 |
| L SPL | 172 | 4.76 | -20 | -76 | 50 |
| L SPL | | 4.48 | -14 | -70 | 44 |
| L LOC | | 3.80 | -26 | -80 | 38 |
| R SPL | 150 | 4.54 | 24 | -60 | 60 |
| R SPL | | 3.28 | 14 | -60 | 66 |
| L preCG | 359 | 4.49 | -8 | -14 | 74 |
| L preCG | | 4.41 | -20 | -16 | 72 |
| L preCG | | 3.99 | -26 | -14 | 60 |
| L SPL | 150 | 4.08 | -28 | -54 | 56 |
| L IPS | | 3.66 | -24 | -60 | 48 |
| L SPL | | 3.11 | -28 | -66 | 58 |
| L Premotor | 96 | 4.00 | -4 | -4 | 46 |
| ***MD > TD*** | | | | | |
| No significant clusters | | | | | |

Abbreviations: FG, Fusiform Gyrus; IPS, Intraparietal Sulcus; IPL, Inferior Parietal Lobules; LOC, Lateral Occipital Cortex; MFG, Middle Frontal Gyrus; MTG, Middle Temporal Gyrus; preCG, Precentral Gyrus; SPL, Superior Parietal Lobule; STG, Superior Temporal Gyrus; L, Left; R, Right. Notes, [†]The left hippocampus subpeaks were identified using AAL masks.

**Table S8.** Brain regions showing significant main effect of time (pre- vs. post- tutoring) on cross-format NRS in Group X Time ANOVA.

| | | | MNI coordinates | | |
|---|---|---|---|---|---|
| **Region** | **Cluster Size** | **T-value** | **x** | **y** | **z** |
| ***Pre > Post*** | | | | | |
| R Frontal Pole | 93 | 3.88 | 28 | 54 | 8 |
| R Frontal Pole | | 3.28 | 30 | 52 | 22 |
| R Frontal Pole | | 3.02 | 30 | 62 | 8 |
| ***Post > Pre*** | | | | | |
| L Hippocampus | 88 | 3.82 | -16 | -40 | 2 |
| L Hippocampus | | 3.28 | -14 | -34 | -6 |
| L Cingulate Gyrus | | 3.26 | -6 | -40 | 0 |
| L Caudate | 161 | 3.79 | -4 | 8 | -2 |
| L Thalamus | | 3.75 | -2 | -6 | 0 |
| R Caudate | | 3.55 | 6 | 22 | 0 |

L, Left; R, Right.

**Table S9.** Brain regions showing significant differences in cross-format NRS between pre- and post- tutoring across all children and in each MD and TD group.

| Group | Region | Cluster Size | T-value | MNI coordinates x | y | z |
|---|---|---|---|---|---|---|
| All children | ***Post > Pre*** | | | | | |
| (MD+TD) | L Hipp | 92 | 3.89 | -16 | -40 | 4 |
| | L PCC | | 3.30 | -6 | -40 | 0 |
| | L PHG | | 3.23 | -14 | -34 | -6 |
| | ***Pre > Post*** | | | | | |
| | R Frontal Pole | 103 | 3.96 | 28 | 54 | 8 |
| | R Frontal Pole | | 3.35 | 30 | 52 | 22 |
| | R Frontal Pole | | 3.07 | 30 | 62 | 8 |
| TD | ***Post > Pre*** | | | | | |
| | No significant clusters | | | | | |
| | ***Pre > Post*** | | | | | |
| | R Putamen | 163 | 5.51 | 28 | 14 | 2 |
| | R Insula | | 4.01 | 28 | 22 | 6 |
| | R Insula | | 3.93 | 34 | 16 | 12 |
| | L IFG | 272 | 5.11 | -42 | 22 | 0 |
| | L Insula | | 4.05 | -28 | 24 | 2 |
| | L Insula | | 3.90 | -42 | 16 | -10 |
| | L Paracingulate Gyrus | 89 | 4.91 | -6 | 46 | -8 |
| | L Paracingulate Gyrus | | 3.65 | -10 | 42 | 0 |
| | L mPFC | | 3.56 | -6 | 54 | -10 |
| | L preCG | 119 | 4.43 | -36 | -8 | 60 |
| | L MFG | | 4.20 | -30 | -4 | 50 |
| | L SFG | | 3.00 | -22 | 0 | 48 |

| | | | | | | |
|---|---|---|---|---|---|---|
| MD | ***Post > Pre*** | | | | | |
| | L Caudate | 585 | 5.66 | -22 | 2 | 16 |
| | L Subcallosal Cortex | | 5.43 | -2 | 6 | -2 |
| | L Caudate | | 5.15 | -8 | 18 | 0 |
| | L Cerebellum | 92 | 4.90 | -26 | -62 | -26 |
| | L Fusiform Gyrus | | 4.07 | -34 | -50 | -20 |
| | ***Pre > Post*** | | | | | |
| | No significant clusters | | | | | |

Abbreviations: Hipp, Hippocampus; IFG, Inferior Frontal Gyrus; MFG, Middle Frontal Gyrus; mPFC, medial prefrontal cortex; PCC, Posterior Cingulate Gyrus; PHG, Parahippocampal Gyrus; preCG, Precentral Gyrus; SFG, Superior Frontal Gyrus; L, Left; R, Right.
